# Translating functional brain activity between humans and monkeys

**DOI:** 10.64898/2026.10.01.755966

**Authors:** K. Braunlich, M. Duyck, S. Duffield, K. Behel, C.I. Baker, R. Conway

**Affiliations:** National Institute of Mental Health; Aix Marseille University; NEI; NIMH

## Abstract

Comparing brain responses across species is essential for identifying models of human cognition and testing theories of brain evolution, but differences in brain size, measurement techniques, and signal dynamics make it challenging. Here we collected dense fMRI data in macaques and humans using the same movies, static-image sequences, and tasks, and developed a multivariate analysis method that aligns neural representations temporally and spatially. The resulting data-driven framework provides a quantitative translation of brain responses among individuals and between species. Using this framework, we uncovered a broadly conserved cross-species hierarchy of retinotopic visual areas and whole-brain correspondence. Yet within this structure, we found that higher-order macaque areas shared more information with lower-order human areas, particularly during movies, providing evidence of species differences. We also identified potential macaque precursors to human regions involved in high-level cognition, including theory-of-mind and reading. These results offer a blueprint for common representational principles across species.

---

Cross-species comparisons are critical for understanding evolutionary origins and evaluating animals as models of the human case (Barron et al., 2020). The macaque brain is a common model of the human brain but is roughly 15 times smaller. To what extent are these species’ brains functionally similar? While convergent evidence supports substantial correspondence for primary visual and motor cortex (Albrecht & Hamilton, 1982; Amunts & Zilles, 2015; Kaas, 2004; Van Essen, 2003; Van Essen et al., 2001), cross-species correspondence beyond these regions is less clear (Mars et al., 2013; Orban et al., 2004). Functional comparisons are particularly challenging because many cognitive tasks are impractical or infeasible in macaques. Macaque fMRI also has lower signal-to-noise ratio, requiring longer scan times (Norman-Haignere et al., 2019). Consequently, functional studies often rely on relatively simple tasks and on statistically powerful univariate analyses (e.g., Lafer-Sousa et al., 2016; Norman-Haignere et al., 2019; Tsao et al., 2008), which can miss functional information expressed in multivariate response patterns or along dimensions not specified a priori (e.g., Aoi et al., 2020; Cromer et al., 2010; Fusi et al., 2016; Mante et al., 2013; Raposo et al., 2014; Rigotti et al., 2013).

To investigate more complex responses, some cross-species studies have exploited naturalistic stimuli, such as movies (e.g., Mantini et al., 2012b, 2012a; Russ et al., 2016). These stimuli sample ecologically relevant feature spaces, elicit strong signal-to-noise responses, and require little behavioral training (Felsen & Dan, 2005; Hasson et al., 2010; Sonkusare et al., 2019). Because movies evoke temporally synchronized responses, they facilitate the use of data-driven multivariate techniques, such as inter-subject correlation (Nastase et al., 2019) and hyperalignment (Fig. 1A-B; Haxby et al., 2011, 2020). Hyperalignment is particularly useful: it maps multivariate responses into a common representational space, allowing shared stimulus-related information to be identified despite individual differences in fine-grained cortical organization.

**Figure 1.**
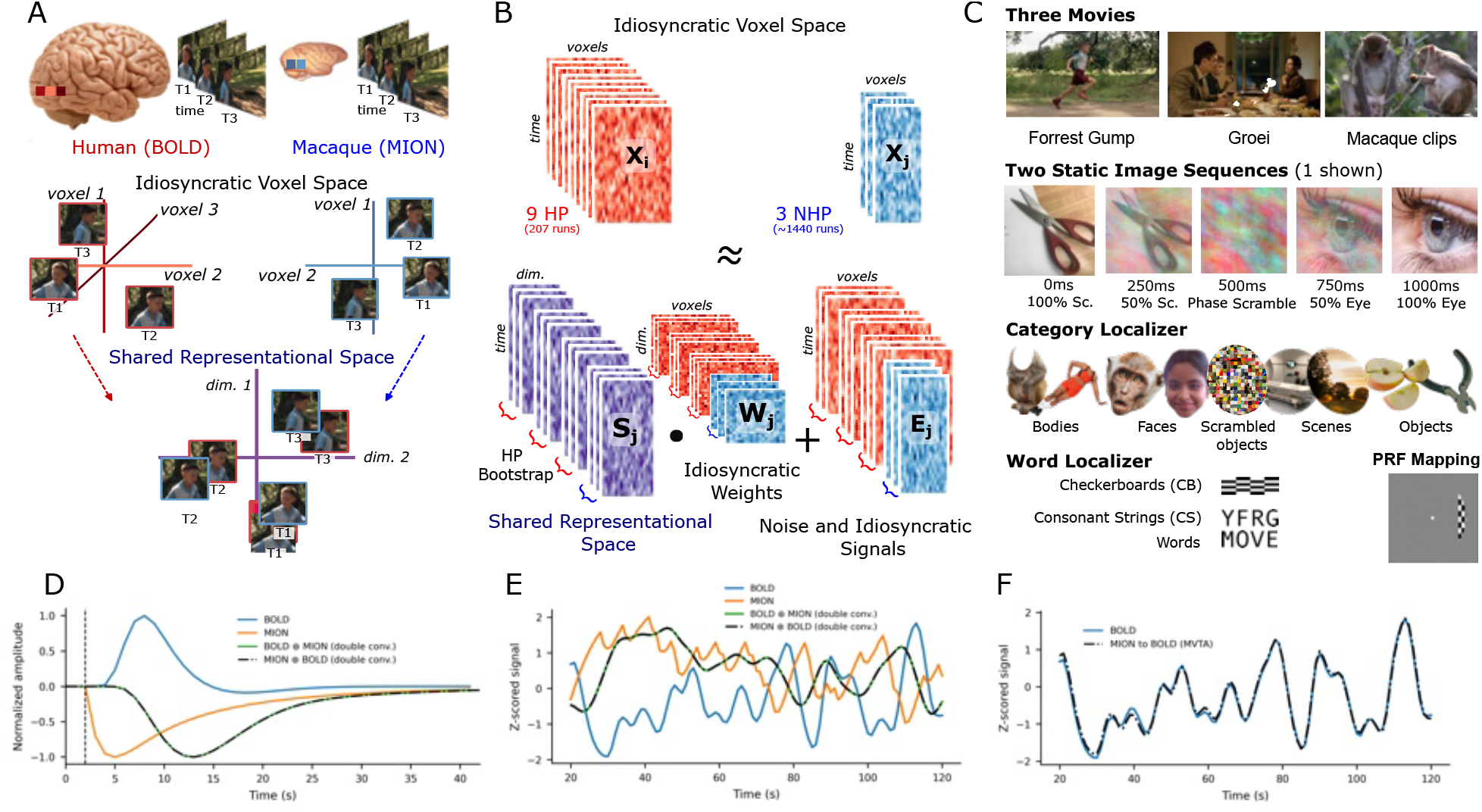
Cross-species spatiotemporal alignment strategy. **(A)** Idiosyncratic voxel organization can misalign shared information. The goal of cross-species hyperalignment is to determine a common representational space into which both species’ data can be transformed. (**B)** Hyperalignment maps each subject’s data X_i_ into a shared representational space S_i_, where dimensions correspond to abstract features rather than voxels. In this example, the shared space has lower dimensionality than the original voxel spaces. Training and testing involved bootstrapped subsampling of humans to match the number of macaques. **(C)** The present dataset comprises responses across five experiments: three movie clips (Forrest Gump, Growth, and macaque behavior clips); two interleaved static-image sequences (1 Hz image presentation); category localizer; word localizer (words, non-words, and checkerboards); and sweeping checkerboard patches for population receptive field (pRF) mapping. **(D)** Canonical BOLD (blue line) and MION (orange line) hemodynamic response functions (HRFs), and results of temporal alignment using the double-convolution method (green: BOLD convolved with MION; black: MION convolved with BOLD; Mantini et al., 2012b, 2012a). **(E)** Simulated BOLD and MION fMRI response to a movie. The double convolution approach aligns BOLD and MION into an intermediate temporal space (dashed black line: MION⊛BOLD, green line: BOLD⊛MION). Although mathematically valid, this approach suppresses high-frequency information and can compound effects of HRF mismodeling. **(F)** Simulated BOLD fMRI response to a movie (blue) and the MION time course after being temporally aligned to BOLD using MVTA (dashed black). MVTA preserved high frequency structure and is fit to individual cross-species ROI pairs, minimizing effects HRF mismodeling.

While hyperalignment holds promise for cross-species comparisons, its assumption of temporally corresponding responses limits its applicability. Basic species differences in temporal responses are further compounded by the use of exogenous contrast agents, such as monocrystalline iron oxide nanoparticles (MION; Leite et al., 2002), commonly used in macaques to boost signal-to-noise (e.g., Norman-Haignere et al., 2019; Vanduffel et al., 2001). MION and BOLD have very different hemodynamic response functions (Fig. 1D).

Here we introduce Spatiotemporal (ST) Hyperalignment, a framework that enables the discovery of shared information across species, accounting for both spatial and temporal misalignment. We apply it to a new fMRI dataset of visual responses collected in macaques and humans (Fig. 1C). We focus on vision because the expansion of visual cortex was likely a driving force in primate brain evolution (Barton, 2026; Kay et al., 2026; Loggia et al., 2025). We present three results.

The first result informs our understanding of the relative organizational structure of retinotopic visual areas V1–V4 within and across species. These areas differ in functional organization (Gattass et al., 2005; Grill-Spector & Malach, 2004; Loggia et al., 2025; Orban, 2008) and are commonly treated as individual entities that correspond across primates. But such correspondence is uncertain beyond V1, and it remains unclear whether these areas constitute a conserved functional hierarchy (Brewer et al., 2005; Fize et al., 2003; Hansen et al., 2007; Kaas, 2020; Kaas & Lyon, 2001; Lyon et al., 1998; Winawer et al., 2010). It is also unknown whether these functional distinctions are reflected in the patterns of stimulus-related information shared across areas during naturalistic viewing. ST-Hyperalignment provides an opportunity to test the degree to which patterns of stimulus-related information among V1-V4 recapitulate a visual hierarchy within each species, whether any organizational structure is conserved across species, and the extent to which putatively corresponding areas align with their counterparts across species.

The second result shows full-brain patterns of cross-species functional correspondence, which allow direct translation of functional brain responses from any anatomical parcel, functional region, or arbitrary collection of voxels in one species to the other species.

The third result demonstrates the utility of the framework as a data-driven discovery tool. We identify potential functional macaque counterparts for human brain areas implicated in complex cognitive functions, including reading (Dehaene & Cohen, 2011; McCandliss et al., 2003) and the ability to attribute mental states to others (theory of mind; Saxe & Kanwisher, 2003).

## RESULTS

Functional MRI data were collected from nine humans and three macaques under matched conditions across five experiments (Fig. 1C): (1) free viewing of movies; (2) fixation-controlled, temporally blended static-image sequences; (3) category localizers defining face-, body-, object-, scene-, and color-selective regions of interest; (4) a word localizer; and (5) drifting checkerboards for population receptive field (pRF) mapping. Movies were expected to recruit widespread cortical networks, whereas static image sequences minimized visual motion and eye-movement effects.

### ST-Hyperalignment: Deriving a common representational space

ST-Hyperalignment is implemented in two stages using data obtained in the first two experiments, which include five datasets: three movies and two static image sequences. Multiview Temporal Alignment (MVTA; see Methods) first learns a convolutional kernel aligning MION data to BOLD (Fig. 1F–G). Then a Shared Response Model (SRM; Chen et al., 2015) learns spatial projection matrices that map each subject’s voxel data into a 50-dimensional common representational space (Fig. 1A–C). Compared to the current approach (Mantini et al., 2012b, 2012a), ST-Hyperalignment preserves high frequency temporal information, and mitigates the potential for HRF mismodeling (Fig. 1D-G; See also SI Discussion and SI Fig. 1).

ST-Hyperalignment was trained and tested with two levels of cross-validation (see Methods). First, by stimulus: the common space was generated using four of the five datasets, repeated five times, holding each dataset out in turn. Second, by individual subject: the left-out dataset was projected into the space and compared across subjects. To quantify the shared information, we divided the held-out dataset into 10 TR stimulus epochs and determined the extent to which the between-subject correlation was higher for matched versus mismatched epochs. We refer to this metric of shared information as stimulus epoch decoding (SED), a variant of an approach used by others (Borovska & De Haas, 2024; Busch et al., 2021; Haxby et al., 2011b; Van Uden et al., 2018).

We used ST-Hyperalignment to assess shared information in ROIs defined with pRF in each species using pRF modeling, cortical atlases, or localizer tasks, as well using unconstrained searchlights. In our first application, we asked whether retinotopic areas V1–V4 show the hypothetical visual hierarchy within each species, and if so, whether this organizational structure is matched across macaques and humans.

### Result 1: The organization of retinotopic visual areas

We consider four hypotheses (Fig. 2B):

- H1 (Unstructured / Supra-Areal): There is no systematic, area-by-area organization of shared functional information (or our approach is insensitive to it), manifesting as unstructured representational correspondence across the regions.
- H2 (Canonical Hierarchy): The assumed canonical functional hierarchy is preserved across macaques and humans, manifesting as maximal correspondence between putatively homologous visual areas and monotonic organization across hierarchical levels.
- H3 (Evolutionary Divergence): Higher-order regions are increasingly specialized within each species, manifesting as increasing functional divergence between successive visual areas (V1 < V2 < V3 < V4).
- H4 (Gaze-Driven Divergence): Early visual areas are disproportionately influenced by species differences in visual behavior (eye movements), manifesting with naturalistic stimuli as decreasing functional divergence between successive visual areas as receptive field sizes increase (V1 > V2 > V3 > V4).

**Figure 2.**
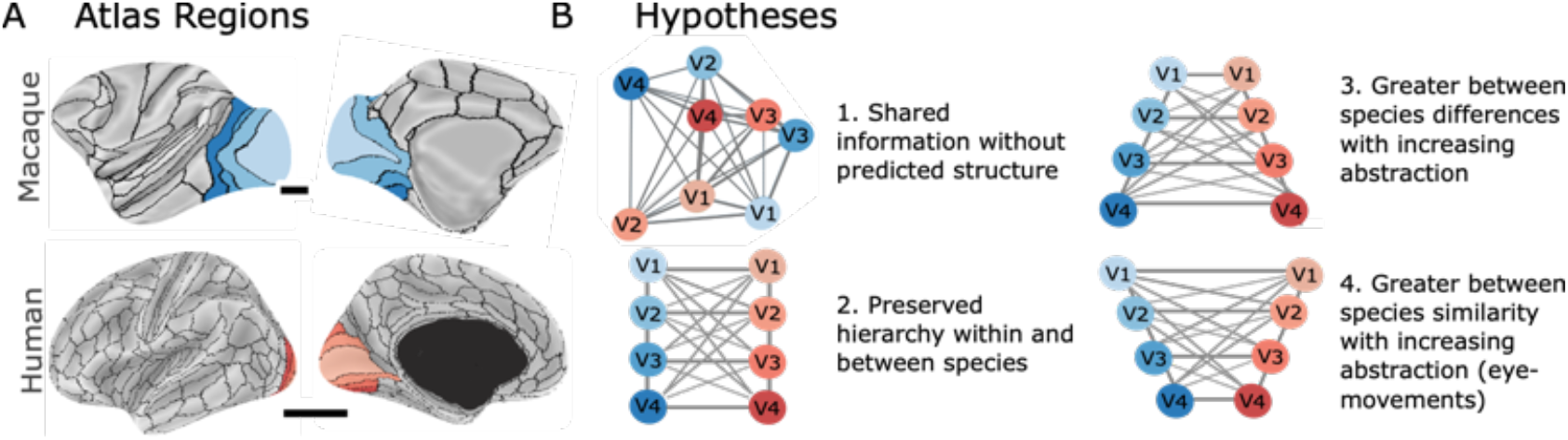
Hypotheses for cross-species representational correspondence in visual cortex. **A)** We focus on areas V1-V4, here illustrated using cortical atlases. Top: Macaque CHARM(4) atlas regions (Jung et al., 2021). Bottom: Human Glasser HCP atlas regions (Glasser et al., 2016). Actual analyses were conducted using individually defined functional population receptive field ROIs.^1^ B) Four hypotheses. **H1**: Unstructured / Supra-Areal. **H2**: Canonical Hierarchy. **H3**: Evolutionary Divergence (Kaas, 2012a, 2020; Rosa & Tweedale, 2005). **H4**: Gaze-Driven Divergence. Human and macaque brains not shown to scale (horizontal black bars indicate relative scale).

We show the within-area and between-area information sharing (SED) as heatmaps. Distinct area-specific representations should produce maximal decoding between putative counterparts, yielding a ridge along the matched-area diagonal.

### Within-species representational organization of retinotopic areas

In both species, shared information decreased with distance along the canonical hierarchy (Fig. 3A–B): for example, V1 shared more information with V2 than with V3. The same pattern appeared across hemispheres, indicating that anatomical proximity alone cannot explain the hierarchy.

**Figure 3.**
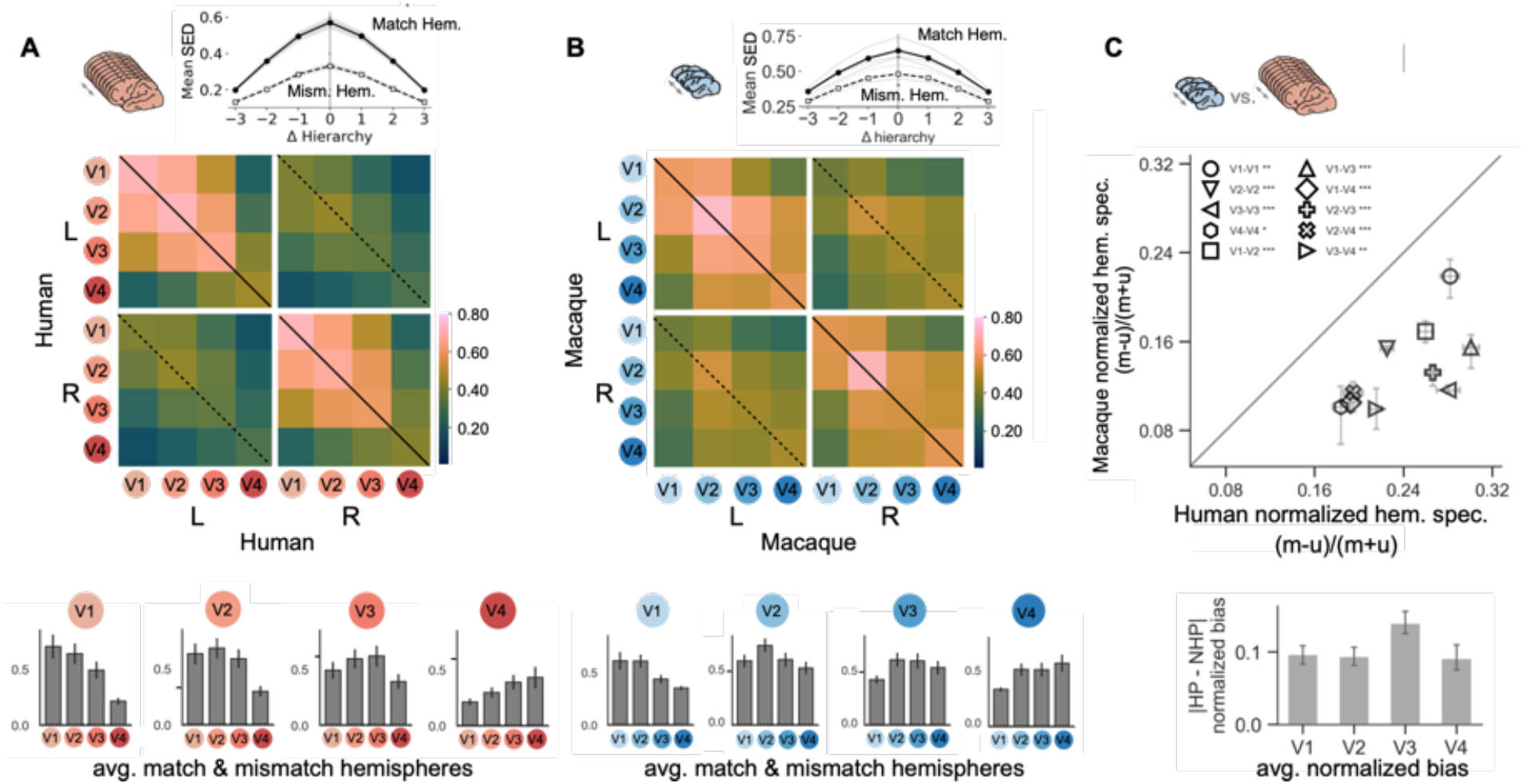
Within species Stimulus Epoch Decoding (SED) results for pRF-derived V1-V4. **(A-B)** Within-species shared information (quantified using SED). Heatmaps show pairwise within-species shared information for all ROI pairs across the left (L) and right (R) hemispheres in humans **(A)** and macaques **(B)**.Warmer colors indicate higher shared information. Chance is less than 1%. The inset plots above each heatmap summarize within species shared information as a function of signed distance in the canonical hierarchy (Δ hierarchy; 0 = same ROI; negative = lower-order target; positive = higher-order target), shown separately for matched (same-hemisphere) and mismatched (cross-hemisphere) pairs. Bar plots show average within-species shared information between the seed (colored disk) and each target ROI, averaged across hemispheres within subject and then across subjects**. (C)** Humans show greater hemispheric specificity than macaques. Values were calculated from normalized within-species SED: (matched hemisphere - unmatched)/(matched + unmatched) from the within-species SED matrices depicted in **A** and **B.** Horizontal error bars denote human SEM, vertical error bars: min./max across macaques; asterisks: FDR-corrected p-values from Welch two-sample t-tests: *** q<0.001,** q<0.01,* q<0.05) . The bar plot at bottom shows the mean difference normalized hemispheric specificity between species (human − macaque) for all pairs involving each ROI.

To quantify the results, we fit a subject-level regression model to each within-species shared-information matrix (see Methods). Shared information was standardized within subject, so a β indicates the direction and relative strength of that effect rather than a raw difference in shared information. A positive area coefficient means same-name pairs (e.g., V1 with V1, V2 with V2) share more information than different-name pairs. A positive hierarchical-proximity coefficient means pairs closer together in the visual hierarchy share more information than pairs farther apart. A positive hemisphere-match coefficient means same-hemisphere pairs share more information than opposite-hemisphere pairs. These three coefficients describe the combined static-image and movie data. Stimulus-type comparisons for within-species effects were not investigated here.

In humans, same-name pairs shared more information than different-name pairs (β = 0.332 ± 0.018 SEM; *t*(8) = 18.32, *p* < 0.001; *W* = 0, *p* < 0.005), and sharing was stronger for areas closer together in the hierarchy (β = 0.492 ± 0.017 SEM; *t*(8) = 28.82, *p* < 0.001; *W* = 0, *p* < 0.005). All three macaques showed the same pattern. Fingerprint coefficients ranged from 0.206 to 0.306, and hierarchical-proximity coefficients ranged from 0.373 to 0.502.

Hemispheric specificity was greater in humans than macaques. Same-hemisphere pairs shared more information than opposite-hemisphere pairs in humans (β = 0.534 ± 0.016 SEM; *t*(8) = 32.99, *p* < 0.001; *W* = 0, *p* <0.005) and every macaque (β range = 0.318–0.427).

To further quantify the hemispheric specificity, each subject’s 8×8 within-species SED matrix was scaled to sum to 1 and reduced to 10 unique unordered ROI pairs (including self-pairs). For each pair, the effect was (matched − mismatched) / (matched + mismatched), and these pair ratios were averaged within each subject. As shown in Fig. 3C, the normalized hemispheric specificity was larger in humans (mean= 0.24, SD=0.012) than in macaques (mean = 0.14, SD = 0.009); roughly a 1.7-fold difference, Welch one-tailed *t* = 14.28, *p* < 0.001; species-label permutation *p* < 0.05, *d* = 8.56. Per-pair tests showed reliable species differences for all ROI pairs after False Discovery Rate (FDR) correction and were strongest for pairs involving V3 (Fig. 3C bar plot).

Overall, within each species, shared information is organized by area and by relative position in the hierarchy. That pattern is inconsistent with H1.

### Cross-species representational organization of retinotopic areas

There is substantial conservation of representational organization across species (Fig. 4A). The highest cross-species SED values remain concentrated near the matched-ROI diagonal, indicating that putatively corresponding visual areas retain similar representational signatures. But the pattern of cross-species SED was modestly shifted toward negative hierarchical offsets, such that greater cross-species SED values can be seen in the lower triangles of the cross-species submatrices than in the upper triangles. The inset line plot in Fig. 4A summarizes this hierarchical asymmetry by averaging cross-species SED values across ROI pairs with the same signed hierarchical offset (Δ hierarchy). A value of 0 corresponds to corresponding areas (e.g., V1–V1), while the magnitude of Δ hierarchy indicates the separation in hierarchical level (e.g., |Δ hierarchy| = 1 corresponds to V1–V2, V2–V3, and V3–V4). Negative values indicate that the macaque ROI is higher-order than the human ROI, whereas positive values indicate that the human ROI is higher-order than the macaque ROI. Cross-species correspondence was shifted such that macaque visual areas aligned more strongly with lower-order human areas than with higher-order human areas.

**Figure 4.**
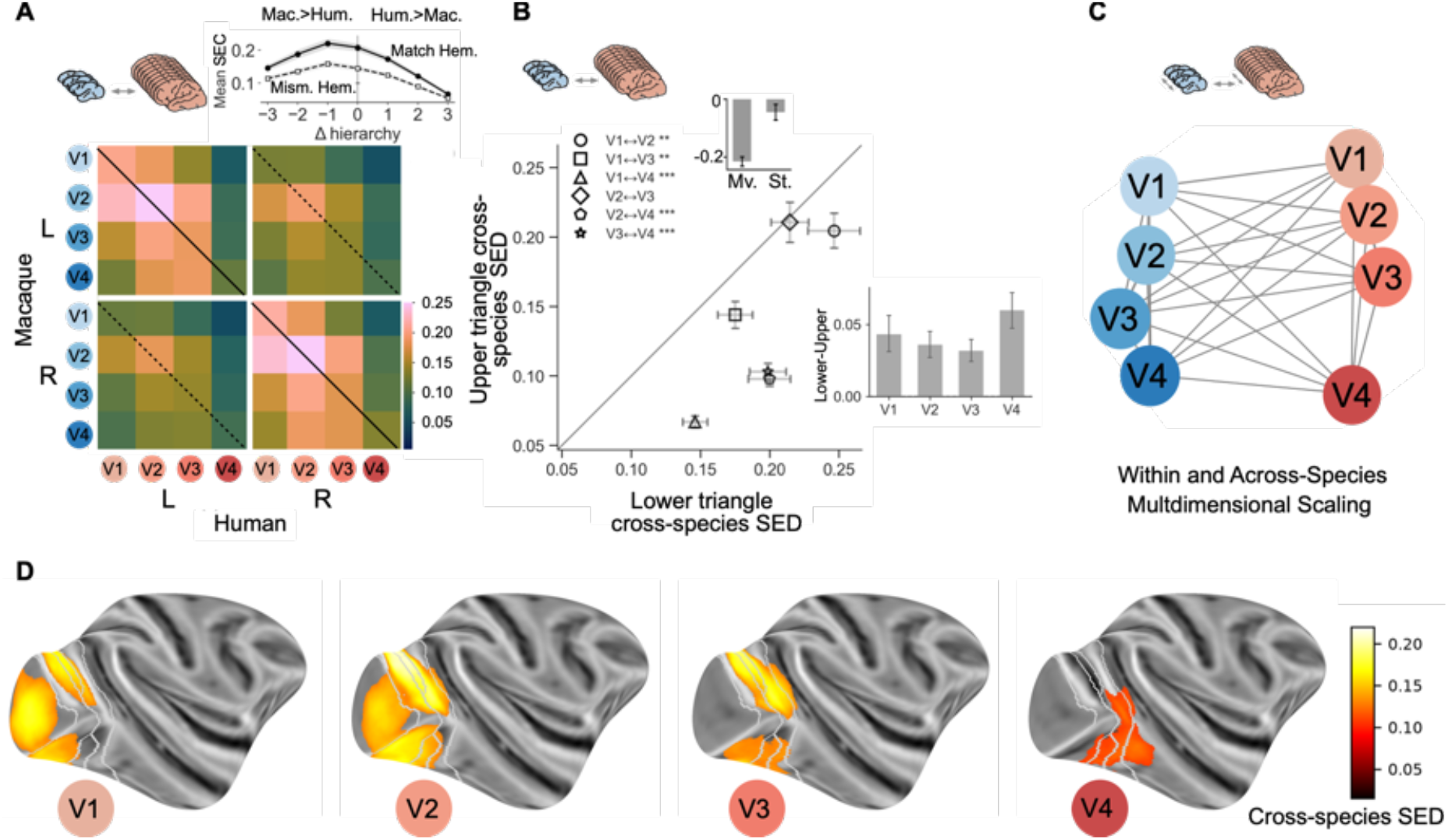
Cross-species Stimulus Epoch Decoding (SED) results for pRF-derived V1-V4. **(A)** Cross-species shared information (SED) between macaque (rows) and human (columns) ROIs. Line plots compare upper- and lower-triangle values as a function o f Δ hierarchy, separately for matched and mismatched hemispheres. **(B)** Higher order macaque brain regions tend to share more information with lower-order human brain regions. Each point represents one off-diagonal ROI pair from A. The x-axis shows lower-triangle SED (macaque > human), and the y-axis shows the corresponding values from the upper-triangle (human > macaque), with the two triangles defined by the main inverse diagonal (solid black line). Values are shown for matched hemispheres. Error bars indicate SEM across subjects; the gray diagonal line denotes identity (*** q<0.001,** q<0.01,* q<0.05). Right bar plot shows mean lower-upper differences (± SEM) for each ROI. Top bar plot shows the matched-hemisphere triangle bias during movies (Mv.) and static images (St.) across V1-V4, computed from balanced-training SED matrices; each bar is the mean across subjects of (upper-lower)/(upper+lower), where negative values indicate greater sharing when the macaque area is higher order than the human. **(C)** Multidimensional scaling using combined within and across species patterns of information sharing recapitulates the putative hierarchy in each species. Distances were calculated from a combined within- and cross-species distance matrix (distance = 1 - SED, averaged across subjects). **(D)** Human pRF-derived seed (right hemisphere V1-V4) to macaque searchlight results.

To evaluate cross-species organization of V1–V4, we fit the same regression framework used for within-species analysis, to the combined stimulus data using default (unbalanced) hyperalignment training (see Methods).

In the full 8×8 regression, coefficients describe effects on within-subject standardized shared information, with other predictors held constant. A positive fingerprint coefficient means same-name human and macaque areas share more information than different-name pairs; a positive hierarchical-proximity coefficient means areas closer in the hierarchy share more than areas farther apart; a positive hemisphere-match coefficient means same-hemisphere pairs share more than opposite-hemisphere pairs; and a positive triangle-bias coefficient means higher-order macaque regions share more information with lower-order human regions than the swapped pairing.

Same-name human and macaque areas shared more information than different-name pairs (β = 0.162 ± 0.010 SEM; *t*(11) = 15.95, *p* < 0.001; *W* = 0, *p* < 0.001), and areas closer in the hierarchy shared more information than areas farther apart (β = 0.317 ± 0.014 SEM; *t*(11) = 22.11, *p* < 0.001; *W* = 0, *p* < 0.001). Same-hemisphere pairs also shared more information than opposite-hemisphere pairs (β =0.293 ± 0.015 SEM; *t*(11) = 19.51, *p* < 0.001; *W* = 0, *p* < 0.001). As with the within-species results, the opposite-hemisphere submatrices in Fig. 4A show the same graded pattern at a lower overall level, demonstrating that the hierarchy is not explained by anatomical proximity alone. Higher-order macaque areas tended to share more information with a lower-order human area than the reverse (β = 0.293 ± 0.026 SEM; *t*(11) = 11.12, *p* < 0.001; *W* = 0, *p* < 0.001). Movies produced higher overall shared information than static images (β = 1.338 ± 0.062 SEM; *t*(11) = 21.49, *p* < 0.001; *W* = 0, *p* < 0.001). Movies also steepened the hierarchical gradient (condition × hierarchy: β = 0.228 ± 0.028 SEM; *t*(11) = 8.26, *p* < 0.001; *W* = 0, *p* < 0.005) and increased the triangle bias (condition × triangle bias: β = 0.400 ± 0.015 SEM; *t*(11) = 26.43, *p* < 0.001; W = 0, p < 0.001). Stimulus type, however, did not reliably change the same-name advantage (condition × fingerprint: β = 0.019 ± 0.019 SEM; *t*(11) = 1.05, *p* = 0.318; *W* = 24, *p* = 0.266) or the same-hemisphere advantage (condition × hemisphere match: β = 0.045 ± 0.021 SEM; *t*(11) = 2.11, *p* = 0.058; *W* = 16*, p* = 0.077). The condition terms were included as effects of no interest, as comparisons between conditions was performed using a balanced training procedure.

Excluding V4,^2^ the triangle bias remained significant but was roughly half as large (β = 0.142 ± 0.028 SEM; *t*(11) = 5.03, *p* < 0.001; *W* = 2, *p* < 0.005). The same regression pattern replicated qualitatively when ST-Hyperalignment was fit under balanced-stimulus training. Balanced training involved learning the hyperalignment *w_i_* on the responses to one movie and one static image sequence in each fold.

To compare triangle bias between movies and static images directly, we used a simpler matched-hemisphere summary on balanced-training shared-information matrices (see Methods). For V1– V3, the lower-triangle bias was reliable for movies (*t*(11) = −6.12, *p* < 0.001; *W* = 0.0, *p* < 0.001) and not for static images (*t*(11) = 1.03, *p* = 0.325; *W* = 26, *p* = 0.34), with movies showing significantly stronger bias than static images (*t*(11) = −4.86, *p* < 0.001). The Fig. 4B inset shows the same analysis for V1–V4: movies (*t*(11) = −13.34, *p* < 0.001; *W* = 0, *p* < 0.001) but not static images (*t*(11) = −1.64, *p* = 0.13; *W* = 16, *p* = 0.08) showed a significant bias, and the difference between conditions was significant (*t*(11) = −7.24, *p* < 0.001).

Because this effect was specific to movies (Fig. 4B, inset bar plot: mean upper minus lower triangle values), it likely does not reflect a fixed difference in the representational properties of these regions across species. Instead, it may reflect species differences in how these regions are engaged or modulated during movies, including differences in attention or sensitivity to visual motion.

A joint multidimensional scaling of within- and cross-species correspondence (Fig. 4C; see Methods) revealed a V1-to-V4 gradient within each species, with corresponding regions from the two species occupying nearby positions. These results most closely support the hypothesis of largely preserved local and global organization between species (H2).

Together, these results reveal a conserved V1–V4 hierarchy alongside two species differences: greater hemispheric specificity in humans and a movie-dependent bias toward stronger correspondence between higher-order macaque and lower-order human areas

### The contribution of eye-movements to representational structure

Although the dominant inter-areal relationships captured by multidimensional scaling (Fig. 4C) do not align with H4 (Gaze-Driven Divergence), gaze may nonetheless contribute to representational structure. We therefore directly tested whether gaze patterns covaried with the neural representations of each ROI during movie viewing using concurrent eye tracking. Differences in scanner eye-tracking configuration, however, affected data yield. Whereas macaque eye tracking was performed with a direct line of sight from outside the bore, human tracking required participants to view the stimulus through a mirror within the head coil. This configuration is more susceptible to pupil-tracking loss, blink artifacts, and calibration drift. As a results, while macaque eye-tracking data were complete, some human data were excluded after quality screening (each human contributed eye-tracking data for at least one movie). This yielded a final dataset of 24 cross-species subject pairs for Forrest, 18 pairs for Growth, and 21 pairs for the macaque movie. Using this dataset, we quantified behavioral and neural similarity: eye-movement similarity (Supplementary Fig. 7) was calculated as the average between-subject Pearson correlation of horizontal and vertical gaze time series, and neural similarity was defined as the Spearman correlation of subjects’ responses within a shared 50-dimensional Shared Response Model space (SRM; Chen et al., 2015).

Across all ROIs, eye-movement similarity was a positive predictor of neural similarity, such that cross-species subject–movie pairings with more similar eye movements exhibited more similar neural response patterns (Fig. 5; see Methods). This effect was strongest in primary visual cortex and weaker in higher visual areas. Explained variance decreased from V1 to V4 (R² = [0.6, 0.51, 0.37, and 0.31], indicating a reduction in the strength of the association between eye-movement and neural similarity across successive visual areas. Coefficients from the Fisher-*z* model showed a generally similar pattern (β = [1.43, 1.19, 0.53, 0.62] for V1–V4, respectively). This is consistent with increasing invariance to location and size, and/or a shift from retinotopic towards conceptual coordinate systems (Pasupathy & Connor, 2001; Roe et al., 2012). Because the observations were not independent (each subject contributed to multiple pairings), we tested these slopes by permuting human and movie eye-similarity labels, with macaque labels held fixed, and refitting the same model. The observed slope exceeded the permutation distribution in every region (all FDR-corrected *q* < 0.05).

**Figure 5.**
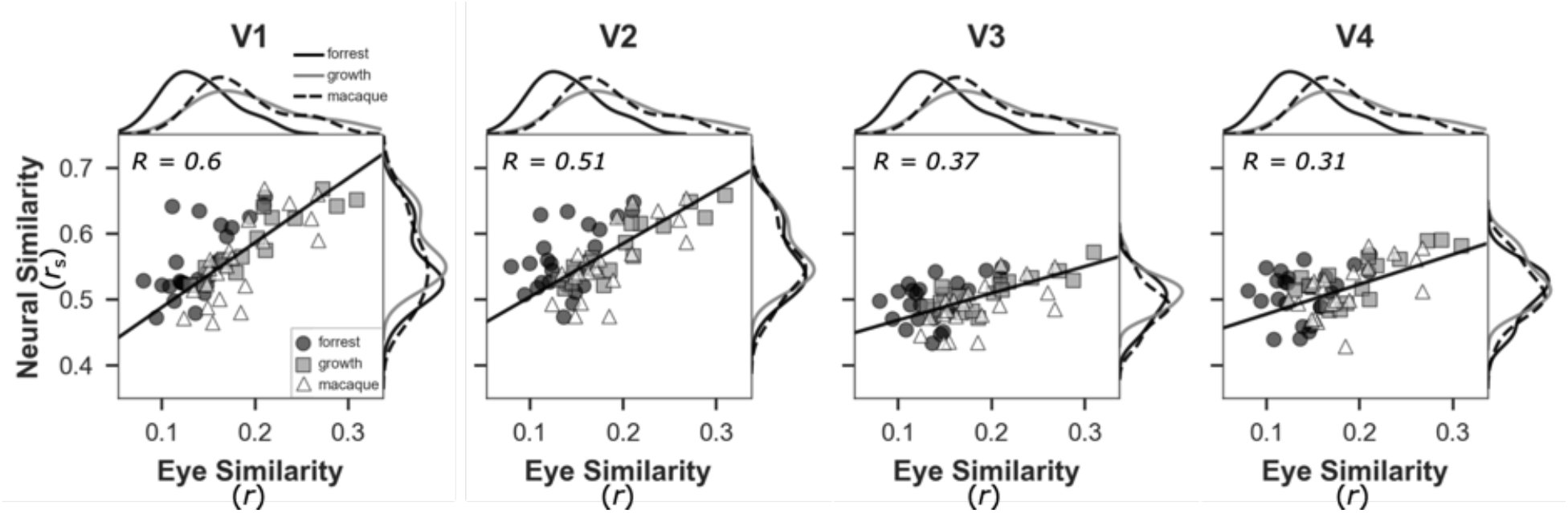
Cross-species relationship between eye-movement similarity and neural similarity in early visual cortex. Scatter plots show eye (Pearson correlation) and neural similarity (Spearman correlation) for each human-macaque subject pair. Marginal distributions depict the corresponding similarity values. The strength of the relationship decreases along the visual hierarchy, with the strongest effects in V1, and weaker, but still significant, effects in V4. Filled circles: Forrest Gump; filled squares: Growth; open triangles: Macaque clips. Black lines indicate least-squares regression slopes in the original units (unstandardized). Positive slopes indicate that subject pairs with more similar eye movements exhibit more similar neural responses.

Taken together, we observed highly structured, area-by-area correspondence across the visual hierarchy, ruling out H1 (Unstructured/Supra-Areal). Gaze similarity predicted neural similarity, with stronger associations in earlier visual areas, consistent with the pattern predicted by H4 (Gaze-Driven Divergence). Gaze did not, however, account for the dominant representational structure. We therefore cautiously interpret our results as supporting H2 (Canonical Hierarchy). The organizational structure was more complex than a simple one-to-one regional mapping, as we observed a significant cross-species hierarchical bias during movies but not static images (Fig. 4B, inset bar plot).

### Result 2: Full-brain patterns of cross-species functional correspondence

The V1–V4 analyses tested a priori hypotheses about patterns of shared information across the visual hierarchy. We next applied the same approach, leveraging whole-brain atlases in each species to explore patterns of information sharing across the cortex. We used the Glasser HCP atlas for human (Glasser et al., 2016) and the CHARM (level 4) atlas for macaque (Jung et al., 2021).To account for regional differences, we learned a distinct HRF transformation for each between-species atlas pair (see Methods).

Using the conservative atlas-wide criterion^3^ (see Methods), the strongest shared information was in occipital and temporal regions (particularly in V1–V4), with considerable cross-species information in parietal and frontal regions of each species (Fig. 6). This matrix was sorted so that strongest SED values were shifted towards the top left corner and organized by lobe and hemisphere.

**Figure 6.**
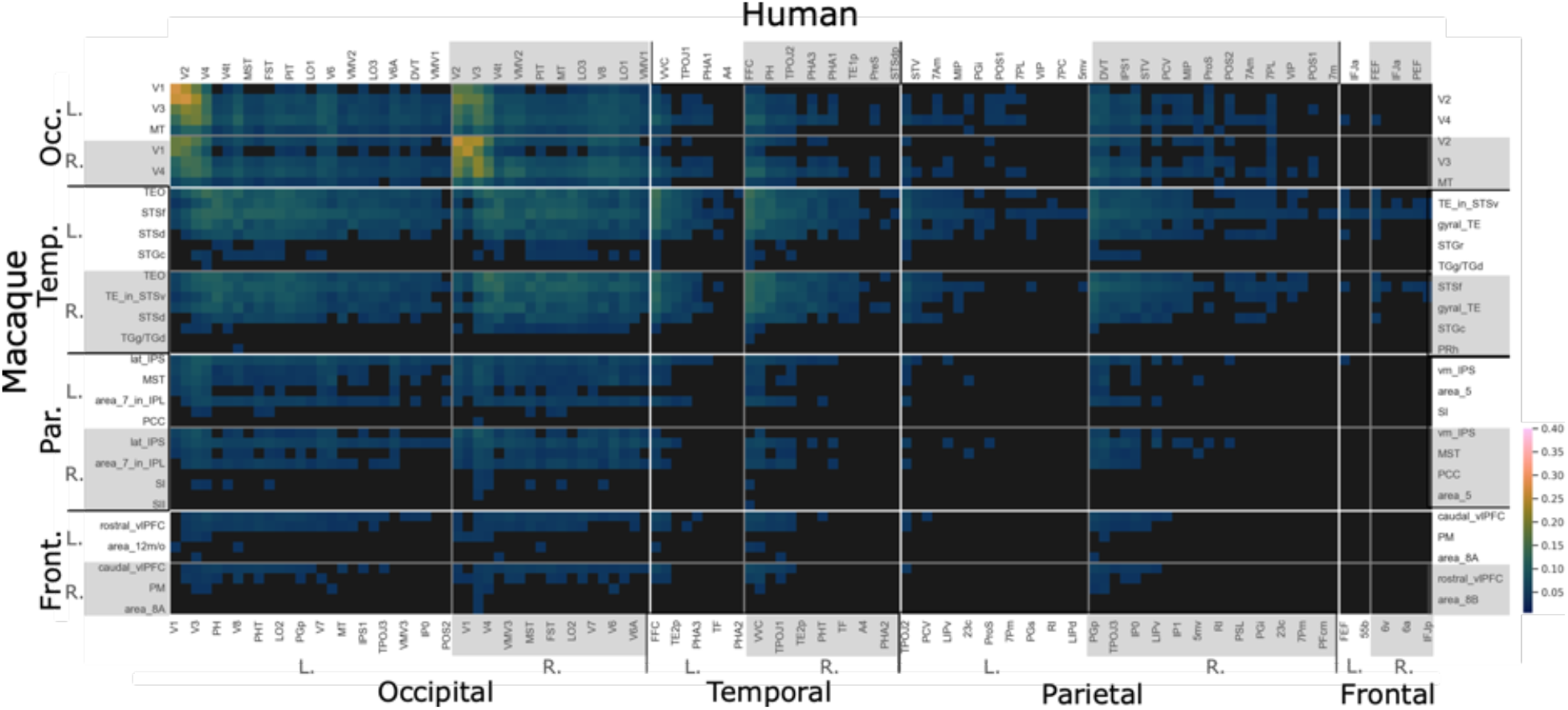
Full brain macaque (CHARM 4) to human (Glasser HCP) atlas cross-species SED accuracy using a highly conservative threshold (all subjects > chance and FDR-corrected mean binomial q < 0.05. Occipital and temporal regions contained the strongest cross-species information, but substantial shared information was also identified in parietal and frontal regions. Overall, the pattern is broadly consistent with conserved sensory processing systems, with higher order regions showing progressively greater species-specific specialization.

To better understand the topology of cross-species organization, we mapped the maximum SED values from each row and column of the matrix onto the cortical surfaces (Fig. 7A). Each macaque ROI was assigned the maximum value from its corresponding row, identifying its strongest human match, whereas each human ROI was assigned the maximum value from its corresponding column, identifying its strongest macaque match. Shared information was widespread across macaque cortex but sparse across large portions of human cortex.

**Figure 7.**
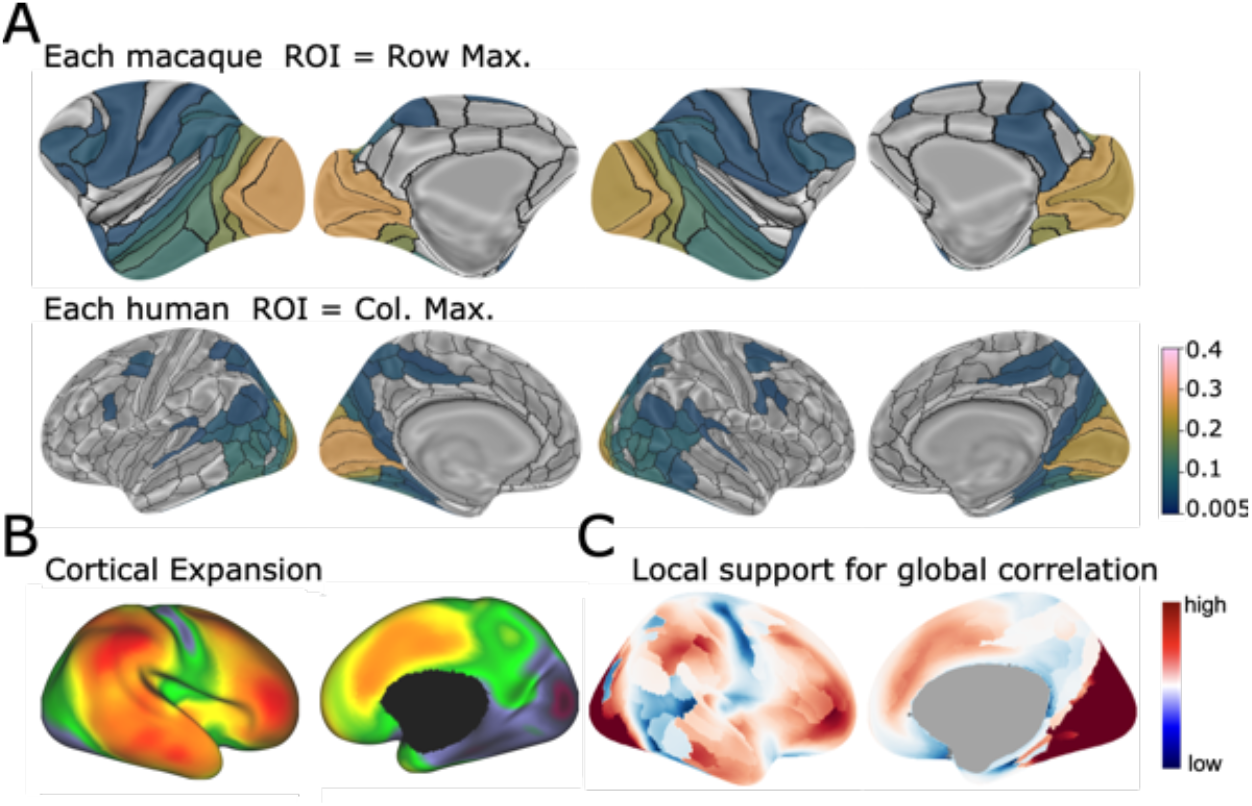
**(A)** Top row: the maximum value of each row of the cross-species matrix depicted in Figure 5 was assigned to the vertices associated with the associated macaque label. Bottom row: The maximum value of each column was assigned to the vertices associated with the associated human label. Although cross-species information was represented widely across the macaque brain, cross-species information in the human brain coverage was sparser, particularly in frontal, parietal, insular, and anterior temporal lobes—areas thought to have undergone disproportionate expansion. **(B**) HCP Cortical expansion map (J. Hill et al., 2010). A spin test indicated a significant but weak correlation (r=0.56, p<0.05) indicating that regions associated with low cortical expansion tended to have stronger cross-species information. **(C)** Local signed support for the full-brain correlation. Red denotes regions where robust shared information colocalizes with low expansion (or weak shared information colocalizes with high expansion). Blue denotes regions that violate this hypothesis. Low support in motor regions is expected due to lack of consistent motor task.

We next compared cross-species information with an independent estimate of cortical expansion derived from the alignment of presumed homologous anatomical and functional landmarks across the human and macaque cortex (Fig. 7B–C; Essen & Dierker, 2007; J. Hill et al., 2010). Across the cortex, regions with less expansion showed greater cross-species information (spin test on sign-flipped SED: r = 0.56, p < 0.05; Fig. 7B; see Methods). The regional map, however, revealed several dissociations from this overall relationship, including regions near the intraparietal sulcus and temporo-parieto-occipital junction that showed both high expansion and high shared information (Fig. 7C). These dissociations highlight the complementary nature of the two approaches: the expansion estimate extrapolates correspondence from presumed homologous landmarks, whereas our approach identifies shared representational structure directly from neural responses during naturalistic viewing. Such dissociations could reflect limitations in the expansion estimate, genuine conservation of function despite substantial expansion, or evolutionary pressure to devote greater cortical territory to functions that remain conserved across species.

### Result 3: Putative macaque counterparts of human-defined functional regions

The analyses presented thus far compared representational structure between pre-defined brain region pairs across species. To move beyond this constraint, we performed cross-species searchlight analyses using human regions as representational seeds and searched the macaque cortical surface for regions exhibiting shared information. Methodologically, this exploratory approach provides an important test of the specificity of the framework. Conceptually, this approach enables the identification of brain regions based purely on shared representational structure. This is valuable for investigating the evolutionary foundations of higher-order cognitive capacities, especially in cases where it is difficult or infeasible to train macaques to perform the same tasks as humans. One motivation for this approach is the hypothesis that novel cognitive functions may build upon and repurpose evolutionarily older representational scaffolds (Dehaene & Cohen, 2007) potentially preserving some representational properties.

First, we applied this approach to the human right temporoparietal junction (rTPJ), which has been associated with theory of mind (Saxe & Kanwisher, 2003). A cross-species searchlight from this seed identified bilateral areas of the macaque superior temporal sulcus (Fig. 8). Activity in this region has previously been associated with social action perception (Ninomiya et al., 2021) and social prediction (Roumazeilles et al., 2021) while its local grey matter density has been linked to social network size (Sallet et al., 2011). These regions also overlap with the proposed macaque counterpart of the human right temporoparietal junction based on connectivity-profile analysis (Mars et al., 2013).

**Figure 8.**
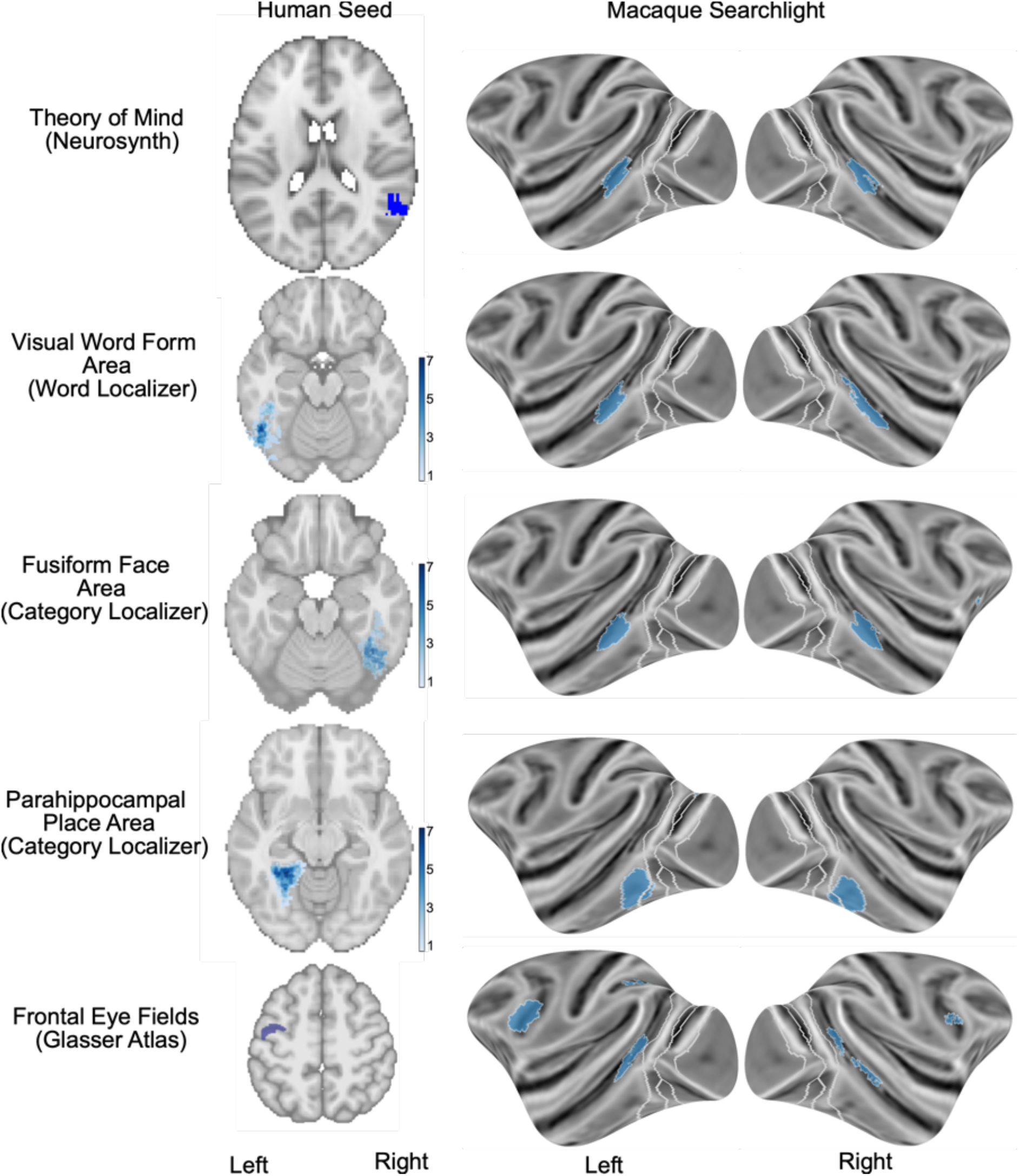
Cross-species searchlight analyses from human seeds associated with higher-order cognitive capacities. This approach is especially relevant in cases where it is difficult or infeasible to train macaques to perform the same tasks as humans. The approach generates data-driven hypotheses for subsequent investigation.

Second, using the functional masks from the human word localizer (see Methods), we identified corresponding bilateral regions in the macaque superior temporal sulcus (STS). These mapped regions closely matched macaque face-selective patches identified via the separate category localizer. This is consistent with prior work showing that word- and face-selective regions occupy similar regions of ventral temporal cortex, potentially reflecting their shared foveal bias (Hasson et al., 2002; Yeatman et al., 2013), with additional evidence that these representations can compete for cortical territory during literacy acquisition (Dehaene-Lambertz et al., 2018; Feng et al., 2022).

Third, using the human fusiform face area (FFA) as a seed, we identified a macaque STS region highly similar to that identified from the human word-selective seed. This convergence is consistent with the overlap between the word-seed mapping and independently localized macaque face-selective cortex.

Fourth, using the human parahippocampal place area (PPA) as a seed, we identified a distinct patch in macaque lateral temporal cortex. The PPA in humans is known to preferentially represent the peripheral visual field (Hasson et al., 2002; Levy et al., 2001, 2004), and the corresponding macaque region likewise shows a peripheral bias (Lafer-Sousa & Conway, 2013), in contrast to the more foveally biased representations associated with words and faces.

Finally, using the human frontal eye field (Glasser et al., 2016) as a seed, we identified shared information in macaque FEF, a foveal region of the STS, and a parietal region associated with visuospatial integration and attention (Pouget, 2015).

## Discussion

In this paper, we introduce a framework, ST-Hyperalignment, which enables a quantitative comparison of multivariate neural representations across species, correcting for differences in brain size and temporal dynamics of measured signals. The framework shifts from asking whether two regions have the same function toward asking where and how representational organization is conserved across the brain. We apply the framework here to macaques and humans, where it revealed a common hierarchical organization of retinotopic visual areas and broader patterns of whole-brain correspondence. Within this conserved structure, we observed a prominent tendency for higher-order macaque areas to share more information with lower-order human areas than expected under identity-mapping, specifically while watching movies, providing evidence of a species difference. Finally, we showcase the power of ST-Hyperalignment as a data-driven discovery tool, using it in a whole-brain search to identify candidate functional counterparts of high-level human brain areas, including those involved in reading (VWFA; Cohen et al., 2000; Dehaene & Cohen, 2011; McCandliss et al., 2003) and in “theory of mind” (rTPJ; Saxe & Kanwisher, 2003; Saxe & Powell, 2006; Saxe & Wexler, 2005; Young et al., 2010).

Cross-species comparisons of brain function have been constrained by three challenges. First, behavioral tasks that probe human cognition are often difficult to match across species, limiting comparisons to simplified stimuli or prespecified stimulus dimensions. Second, hemodynamic signals differ across species and acquisition methods, obscuring shared neural dynamics and introducing systematic temporal biases. Third, corresponding brain regions across species may contain a mixture of neural signals that are shared versus species-specific (Anderson, 2007, 2010; Dehaene & Cohen, 2007; Kaas, 2012b; Krubitzer, 2007; Krubitzer & Hunt, 2007; Krubitzer & Kaas, 2005), making the conserved component difficult to isolate. ST-Hyperalignment addresses these challenges by using multiview temporal and spatial alignment to target shared cross-species structure, and between-species decoding to identify shared information. Naturalistic movies and interleaved static images provide a common stimulus space across species; MVTA accounts for differences in the hemodynamic response; and hyperalignment identifies shared multivariate structure without requiring voxel-wise correspondence. Below, we discuss the key findings afforded by this approach.

We first applied ST-Hyperalignment using data obtained during naturalistic vision to test the extent to which retinotopic visual areas constitute a hierarchy, and one that is conserved across macaques and humans. Across V1–V4, putatively corresponding areas shared more information than mismatched areas, and the resulting representational fingerprints were reproducible across individuals within each species and conserved across species. Multidimensional scaling of the dominant representational axes revealed a consistent hierarchical V1–V4 organization in both species. Information sharing declined with distance along this sequence, both within and across hemispheres, showing this organization cannot be explained by anatomical proximity or spatial smoothing. Together, these results support the hypothesis that V1-V4 are organized in a hierarchy, and that this hierarchy is conserved across macaques and humans. These findings imply that electrophysiological observations made in the macaque apply to the human case and have implications for accounts of primate brain evolution.

But within this shared hierarchical organization, ST-Hyperalignment uncovered an unexpected cross-species difference: putatively higher-order macaque visual areas were biased to share more information with putatively lower-order human visual areas (Fig. 4B). This result was stronger for data acquired during movie watching than during static image sequences, suggesting that it does not reflect an intrinsic difference in areal properties. Instead, this hierarchical shift is consistent with species differences in behavioral goals during naturalistic vision. Such differences could manifest as different contextual understanding, attentional modulation, and motion signals from higher-order networks that differentially modulate early visual representations (Gilbert & Li, 2013; Roelfsema & De Lange, 2016).

Differences in behavioral goals are classically reflected in eye-movement patterns (Hayhoe & Ballard, 2005; Rothkopf et al., 2016; Yarbus, 1967), and prior work has shown that humans and macaques show both shared and species-specific gaze patterns during naturalistic viewing (Shepherd et al., 2010). We therefore investigated the relationship between eye-movements and cross-species neural information sharing. We found that cross-species subject pairs with more similar gaze patterns showed more similar neural representations, an effect that decreased progressively from V1 to V4. This pattern of results is consistent with a hypothesis in which eye movements and behavioral goals are linked, and the linking has a greater impact on visual areas with smaller receptive fields and lower tolerance to spatial jitter (Borovska & De Haas, 2024; Rust & DiCarlo, 2010). The impact of gaze dissimilarity demonstrates that unconstrained behavioral variables impart distinct structure that should be considered in cross-species analyses. Nevertheless, this effect remains secondary to the conserved hierarchical structure that organizes V1 through V4.

Beyond visual cortex, we used ST-Hyperalignment to determine broader patterns of cross-species correspondence across the brain. These analyses confirmed strong information sharing in occipital cortex, as expected from the analyses of V1 to V4, but also uncovered substantial information sharing in temporal, parietal, and frontal regions. These results show that visual stimuli evoke activity extending beyond vision to include cognition. But notably, while most of macaque cortex contained shared information with human, many human regions did not share information with the macaque—in particular, frontal cortex in humans had very low information sharing with macaque, providing possible evidence of the evolution of human-specific cortical areas.

Across the cortex, shared information tended to be greatest in regions with low putative evolutionary expansion. One notable exception was motor cortex, which showed low information sharing despite relatively low expansion, likely reflecting the absence of a controlled motor task in the movie paradigm. Conversely, several regions classified as having high putative evolutionary expansion showed substantial shared information. This pattern is notable because greater evolutionary expansion is often taken to reflect increased cross-species divergence in cortical organization and, consequently, might be expected to limit the correspondence of functional representations across species.

ST-Hyperalignment revealed substantial sharing in highly expanded temporoparietal (TPOJ1–3), lateral temporal (PHT, TE1p, TE2p), and intraparietal association cortex (LIPv, IP1), as well as motion-sensitive visual cortex (MST; Supplementary Table 1). The finding is particularly notable in temporoparietal and intraparietal cortex, which support functions including attentional control, visuospatial processing, social cognition, and theory of mind (Corbetta & Shulman, 2002; Igelström & Graziano, 2017). Thus, regions estimated to have undergone substantial evolutionary expansion do not uniformly show reduced functional correspondence across species. Instead, substantial cross-species information sharing in some highly expanded regions may indicate that conserved functional organization can be maintained despite large anatomical changes. One possibility is that these regions experienced selective pressure to preserve important computations during cortical expansion; another is that expansion provided additional representational capacity while retaining aspects of an ancestral functional organization.

Finally, we used ST-Hyperalignment as a data-driven tool to discover functional counterparts in macaque of regions associated with high-level cognitive functions in humans. The “Theory of Mind” region in human right temporoparietal junction (TPJ) shared the most information with macaque superior temporal sulcus (STS), bilaterally. Interestingly, these putative functional corresponding areas overlap with regions previously implicated in social processing (Ninomiya et al., 2021; Roumazeilles et al., 2021; Sallet et al., 2011) and with areas proposed as connectivity-based counterparts of human TPJ (Mars et al., 2013). Meanwhile, the visual word form area (VWFA), which has been associated with reading, shared most information with bilateral macaque regions in foveally biased regions of extrastriate IT cortex. These putative functional corresponding regions closely overlap with results from human fusiform face area (FFA) and were different from those associated with the human parahippocampal place area (PPA). The PPA shows a peripheral bias (Epstein & Baker, 2019; Hasson et al., 2002; Silson et al., 2015), as does the region in macaques we identified using ST-Hyperalignment (Arcaro & Livingstone, 2017; Lafer-Sousa & Conway, 2013).

Shared representational structure may reflect conserved computational building blocks without implying identical high-level functions. For example, representational overlap between human TPJ and macaque STS may reflect conserved mechanisms for processing dynamic social cues without implying equivalent theory-of-mind reasoning. Similarly, culturally novel capacities like reading do not require de novo cortical structures, but instead repurpose evolutionary precursors—namely, ancestral ventral visual circuits optimized for complex feature recognition (Dehaene & Cohen, 2007). Cross-species mappings can therefore nominate candidate functional counterparts and generate testable hypotheses about which aspects of representational architecture have been preserved across lineages.

Importantly, conservation of representational organization does not require conservation of the behavior or higher-level computation that ultimately depends on it. A human-specialized function, for example, may rely on a circuit whose lower-level representational structure is partly conserved in macaques. The value of a data-driven cross-species approach is therefore not to assign human functional labels to macaque cortex, but to identify where and to what extent representational organization is conserved. Targeted experiments, including electrophysiology and stimuli or tasks designed to isolate specific candidate computations, can then determine the nature of that correspondence. Future work could also use ST-hyperalignment to test other systems proposed to be functionally conserved across primates, including networks involved in face and color processing.

More broadly, ST-Hyperalignment provides a framework for extending comparative functional mapping beyond the human–macaque comparison to other species and subject populations. MVTA may also be useful independently for aligning signals obscured by unknown temporal filters across imaging modalities, species, and populations. Because these analyses produce rich whole-brain maps that cannot be fully captured in a conventional paper, we have made our searchlight results available through an interactive website for further exploration. Together, these tools provide a flexible framework for comparing brain function across biological and methodological boundaries, and for charting both conserved and divergent representational organization across species.

## METHODS

We collected fMRI data across 3 experimental paradigms (Fig. 1C). This included naturalistic stimuli (movies and interleaved static image sequences), block-design localizers, and population receptive field (pRF) mapping (Dumoulin & Wandell, 2008a). Following previous work (Norman-Haignere et al., 2019), we collected roughly 20 times more data for macaque than for human to obtain similar SNR.

### Subjects

#### Human

Twelve human participants completed all experimental paradigms (8 female, age: M = 26, range = 22-31 years old). All were screened to have normal or corrected-to-normal visual acuity, normal color vision, be right-handed, and have no history of psychological or neurological issues. Participants provided informed consent in participation and data sharing and received financial compensation for participating in the studies. The NIH Institutional Review Board approved the research as part of study protocols 93-M-0170 (NCT00001360) and 18-EI-0046. Data from three of these participants were excluded from analysis because of excessive head motion (framewise displacement greater than 1.2mm), leaving nine participants in the analyzed sample.

### Macaque

Data was collected on three male rhesus macaque monkeys (Macaca mulatta), Ri (∼8.5 kg, 7.5 yo), Mo (∼11 kg, 7.5 yo) and Wo (9.12 kg, 9 yo). All experimental procedures and animal care were approved by the National Eye Institute Animal Care and Use Committee and complied with the United States Public Health Service Policy on Humane Care and Use of Laboratory Animals. MR-compatible plastic head-posts were surgically implanted on each animal under isoflurane anesthesia in aseptic conditions. Two of the three animals (Ri and Mo) had participated in any previous studies, but each had been trained to receive a reward for fixating a dot (see Supplementary Materials for training information).

### Stimuli & Tasks

#### Movies

We used three naturalistic movie clips. “Growth” (Dutch: "Groei"; van der Woerd, 2016) follows a family across many years but is edited to create the appearance of a continuous single take, without abrupt cuts. We also used a segment from “Forrest Gump” (Zemeckis, 1994; the commercially available 1080p DVD release). The third movie was a collection of clips of macaque monkeys in the sourced from YouTube. All three videos lasted 530 sec. and were presented without sound. The size of the videos in degrees of visual angle was 22.0° by 9.2° for Growth, 22.0° by 9.3° for Forrest, and 18.0° by 10.1° for Macaque. Each run started with 5 TRs showing the movie’s first frame and ended with 10 TRs (8 TRs showing the movie’s last frame and then just the fixation cross). Each movie run therefore totaled 227 TRs (about 9.5 minutes).

### Static Image Sequences

To reduce sources of variability associated with movie viewing, including eye movements and visual motion, we also collected data while subjects viewed temporally synchronized static-image sequences lasting 168 TRs (7 min each). Successive intact images were presented at a rate of 1 Hz. To reduce neural responses to abrupt stimulus onsets and offsets, for each pair of adjacent intact images we generated a single fully phase-scrambled intermediate image derived from the two images. The displayed sequence then smoothly interpolated from the first intact image to this phase-scrambled intermediate and from the intermediate to the next intact image, with each 1-s interval comprising 20 frames of 50 ms each. Each frame subtended 12° × 12° of visual angle. All participants except one macaque performed a one-back task, with three stimulus repetitions occurring at random times per minute.

Each of the two static image sequences involved the randomized presentation of 400 unique images drawn from 6 visual categories (1/8 animals, 1/8 bodies/body parts, 1/8 faces, 1/8 foods, 1/4 objects, and 1/4 scenes). The objects included tools, food, and other items linked to hand-related affordances. Half of the scene stimuli depicted indoor environments, while the other half featured natural outdoor settings. For the body and face categories, half of the images were of humans and half of monkeys. Similarly, for food, objects, and indoor scenes, half of the stimuli were chosen to be familiar to the monkeys (e.g. objects within the lab, lab rooms and hallways).

Each run started with 5 TRs showing the first frame of the video and ended with 10 TRs (8 showing the last frame and 2 of fixation cross), totaling 183 TRs in all. Participants were asked to maintain fixation on a white central fixation cross (0.5° wide).

All subjects were shown two static image sequences. For humans, each sequence was presented twice. For monkeys, because of the high number of repeats of these two sequences, we interleaved novel sequences (constructed from the same 800 images with the same method) that were not included in analyses.

### Block Design Localizers

Participants viewed two types of block-design localizer tasks. The category localizer included 1440 unique images with stimulus blocks for faces, bodies, scenes, objects, and scrambled objects. The word localizer included three types of stimulus blocks: words, consonant-strings, and checkerboards (which were matched in size and screen location).

In both localizer tasks, conditions were presented following an “ABN” design (Maus et al., 2010) such that 4 null blocks (consisting of grey screen with fixation) were included in each run. Each block lasted 15 s and comprised 24 stimulus presentations (4 trials per 2.5s TR). Each stimulus within each block was presented for 525 ms, followed by a blank inter-stimulus interval of 100 ms. All subjects (except one macaque) performed a one-back recognition task (matching exact stimulus identity) while maintaining fixation on a target presented at the center of the screen. The first block was preceded by two blank screen TRs (fixation cross on grey screen) and concluded with 10 TRs with the same blank screen.

The four separate blocks of each category within a run contained 4 or 5 one-back matches per block (the order was randomized). This choice was made to reduce the predictability of the task while holding the number of 1-back matches the same for all conditions within a run. The order of conditions within the runs of each type (including the null condition) was generated using a balanced Latin squares design. Across runs, each condition was preceded by every other condition an equal number of times. Within each run, the same condition order was repeated four times.

The category localizer included 1440 unique images with stimulus blocks for faces, bodies, scenes, objects, and scrambled objects. We used 288 unique images for each of these 5 categories. Half of the faces were human faces taken from the FFHQ dataset (Karras et al., 2019), with primarily neutral facial expressions but with variations in ethnicity, age, viewpoint, and gender. The other half were monkeys (mostly macaques) with similar variations from the PrimFace database (https://visiome.neuroinf.jp/primface). For (headless-) bodies, half of the images were from humans and half were from monkeys. For scenes, half the scenes were indoor scenes (half were photos taken from the animal environment), and half were outdoor scenes (half natural scenes and half human-made). The object domain consisted of sub-domains: 1/3 foods, 1/3 hand or mouth-held object that would fit within a roughly a two square foot box, and 1/3 larger objects. The objects were chosen so that most object classes were familiar to the animals.

Finally, scrambled object images were created by first extracting boxes from the individual object images (which were 1000 x 1000 pixels). This involved dividing each image into a set of 20 x 20 grids (400 grid locations total). To discard mostly empty grid locations, boxes with a mean alpha channel (a binary mask) value of less than 0.3 were then discarded. This process was used to create a set of boxes for each object image. Once a set of object boxes was extracted from each image, a novel image (1000 x 1000 pixels divided into a 20 x 20 grid) was constructed by pulling random boxes from this set. Each scrambled object image comprised small boxes sampled from many different images. This choice maximized variability within each scrambled image and accentuated the vertical and horizontal lines defining the grid boundaries. To further minimize adaption to these grid boundaries, the phase of the scrambled images was shifted vertically and horizontally in 20 steps of 1 pixel (i.e., the size of a single box).

Faces, bodies, and objects were cropped from their backgrounds. To similarly introduce variation in the spatial profile of the scrambled-object and scene stimuli, these images were presented within elliptical apertures that varied in orientation and aspect ratio. Within each run, ellipse orientation ranged from 0° to 180° in 96 steps of 1.875°, and aspect ratio ranged from 0.6 to 1.0 in six steps of 0.08. Each orientation and aspect ratio occurred equally often within a run, and the same sequence of aspect ratios was used for the scrambled-object and scene categories. Across the six runs, the correspondence between orientation and aspect ratio was shifted by one step on each run, such that, across runs within a session, each orientation was paired once with each aspect ratio. These apertures were then randomly assigned to the unique images in the scene and scrambled-object categories.

One-back repeats involved presenting the same scene or scrambled object within the same aperture. All stimuli were within a 1000 px side square presented at a size of 12°, and their position was jittered within a 0.5° window (note that cropped stimuli did not span the whole 1000 px and had a mean height ∼8.5° and width of ∼7.8°). Each block type could be presented in full color or greyscale. Greyscale versions of each image were generated separately for each setup by converting RGB values to CIE xyY coordinates using monitor-specific spectral measurements, replacing the x and y chromaticity coordinates with those of the grey background while preserving luminance (Y), and converting the resulting values back to RGB. Humans performed 6 runs in a single session. Each run comprised 24 blocks and lasted 156 TRs.

The word localizer included three types of stimulus blocks: words (four letters), consonant-strings and checkerboards. Consonant strings were matched to a four-letter word by replacing all vowels of the word with a consonant and shuffling the final letters. Both words and consonant strings were presented in uppercase letters of the Roman alphabet at a width of 3°. Checkerboard boundaries matched the size of the lettered stimuli and consisted of black and white rectangles. These rectangles varied in size randomly between 1.5 and 11 cycles along the horizontal axis and between 1 and 3.5 along the vertical axis. The center of the stimulus was jittered within a 0.3° window from trial to trial, and fixation was a 0.25° diameter red dot.

### Population receptive field mapping

We used a conventional population receptive field mapping stimulus consisting of a moving bar that traversed the screen in eight different directions within a circular aperture of the maximal visible diameter of the human setup (15.0°). The moving bar had a width of 15% of the aperture diameter. It contained a full contrast checkerboard pattern whose components (0.25° wide) were moving at 4 degrees visual angle (dva) per second alternating between movement parallel (one period back and forth) followed by orthogonal to the bar (one period back and forth). The bar sweep spanned the full diameter in 14 equally spaced steps. Each bar position was present on the screen for 1 TR (2 s); therefore, one full bar sweep lasted 28 s. The direction of sweeps was Left-Right, BR-TL, Top-Bottom, BL-TR, R-L, TL-BR, B-T, and TR-BL, and was kept constant across runs. Each direction was presented once in a run. Two blank TRs preceded the onset of the first bar, and ten blank TRs followed the offset of the last bar, totaling 124 TRs in a run.

Participants were instructed to maintain fixation on a central cross of 0.3° and press a key whenever they detected a change in the cross luminance. These changes occurred for three frames, were uniformly distributed every three to ten seconds, and were chosen at random from four constant levels of difficulty in each direction (luminance increment or decrement). Correct responses were defined as button presses within two seconds following change onset. For humans, we collected ten runs in the retinotopy session.

We defined V1-V3 using standard criteria. We defined human V4 as described by Winawer et Witthoft (2015).

### Procedures & Apparatus

#### Human procedures

For humans, each session started with a reminder of the instructions regarding minimizing motion and following the dot during the gaze calibration procedure, followed by verbal instructions specific to the runs for that session. Each run usually started with a 9-point calibration process followed by written instructions that ended with directions to look at the fixation dot/cross when ready to start (thus ensuring correct screen mirroring). Whenever the run contained a behavioral task, and participants were given the response box in their right hand, they were asked to choose a finger and a button to use and to maintain consistency using these during the session. Feedback on task performance was displayed at the end of each run when relevant.

### Macaque procedures

Before a scanning session, MION contrast agent (Predescu et al., 2018) was injected into the saphenous vein of the animal to increase the signal-to-noise ratio (Vanduffel et al., 2001). MION doses were titrated for each subject during the first few sessions to reach a reduction of ∼50/60% signal intensity from EPI BOLD signal intensity.

Macaques were rewarded during the session with drops of water or juice for performing the different tasks of each paradigm. During free viewing (movies), monkeys were rewarded at regular intervals when their gaze was within the video aperture. During fixation (temporally synchronize static images, domain localizers, and retinotopy), they were rewarded with a small reward at regular intervals for maintaining their gaze in a circular window of usually 1° radius. A bigger reward was given when giving a correct response to the task. Reward interval and quantity were adjusted during the session to adapt to the animal’s motivational state.

### Human apparatus

Stimuli were presented on a 32-inch LCD monitor at the end of the MRI bore (1920 × 1080 pixels, 120 Hz - Cambridge Research Systems Ltd.), at 151.5 cm, subtending 26.5° × 15°. Participants viewed the screen through front surface mirror glasses (attached to the RF coil). An EyeLink 1000 Plus infrared eye-tracking system (SR Research Ltd.; Pupil-CR mode, 1000 Hz sampling rate) was attached to the bed behind the coil and used to track the right eye. A pulse oximeter was placed on the left index finger. Respiration and cardiac activity were recorded at 1000 Hz (MP150, BIOPAC Systems, Inc.) but were not used in analyses. Participants made behavioral responses with fingers of their right hand (Lumina LU-400, Cambridge Research Systems Ltd.). Stimuli were presented using the Shady Python module (N. J. Hill et al., 2019).

### Macaque apparatus

Monkeys sat head-fixed in the sphinx position in a horizontal MR-compatible primate chair (Rogue Research Inc., 2026). Stimuli were presented on a rear projection screen via a PROPixx projector (1920 × 1080 pixels, 120 Hz - VPixx Technologies Inc.). The screen was 61.5 cm from the eyes, subtending 34.5° × 19.8°. One eye (usually the right) was monitored using an Eyelink 1000 Plus (SR Research Ltd.) at 1000 Hz. Custom-made lever(s) signaled behavioral responses (right lever press for M1, both levers release for M2) through optic fiber light reflection (https://github.com/NEI-LSR/MRI-ResponseBox). Liquid was delivered using a pressure-based liquid-delivery system (Mitz, 2005). A DATApixx hub (VPixx Technologies Inc.) was used for communication between stimulus, reward, lever, and eye-tracking systems.

Although the two species were scanned in different setups with different display sizes, we maximized the similarities across setups. Notably, the size in degrees of visual angle of all stimuli and the refresh rate were identical; both displays’ red, green, and blue channels were quasi-linear, as measured with a PR-655 Spectroradiometer (Photo Research Inc.), and the maximum luminance was between 110 and 120 cd·m⁻².

### fMRI Acquisition

#### Human fMRI acquisition

Human MRI data were collected using a 7T Siemens Magnetom Terra, with a Nova 32-channel RF coil at the Clinical Research Center on the National Institutes of Health main campus. Each participant performed 3-4 sessions, each of which lasted between 1.5 and 2 hours. We collected several whole-brain anatomical measures, including a T1-weighted MP2RAGE sequence (0.75 mm isotropic resolution, TR = 4.3 s, TE = 1.99 ms, TI1/TI2 = 840 ms/2370 ms, FA1/FA2 = 5/6°), a T2 sequence (voxel size: 0.5 x 0.5 x 2 mm, TR = 2.68 s, TE = 80 ms, FA = 100°) and a FLAIR sequence (1 mm isotropic resolution, TR = 9 s, TE = 269 ms, TI = 2.6 s, FA = 120°). We also collected a whole brain T1-weighted MPRAGE sequence at the beginning of each session to help with alignment and slice placement for functional data collection (0.75 mm isotropic resolution, TR = 2.3 s, TE = 1.75 ms, TI = 1.05 ms, FA = 6°). Functional imaging was collected using T2^* weighted echo-planar imaging (EPI) sequences (1.2 mm isotropic, TE = 23 ms, FA = 55°, partial Fourier 6/8, iPAT 2, multiband (MB) 2). Repetition times and acquired slices varied across the different experiments. For retinotopy, we used a 2 s TR (TR = 2 s, 44 slices, no gap between slices), whereas, for all other experiments, we used a 2.5 s TR (TR = 2.5 s, 76 slices, 10% slice gap, i.e. 0.12 mm). For retinotopy, slices were positioned to optimize visual and temporal coverage and at an angle with the calcarine sulcus, and for all other experiments, most of the brain was covered (Supplementary Fig. 6). All runs were collected with the Anterior-to-Posterior phase encoding direction. At the end of each run, while the participant was still fixating on the blank screen, we collected several TRs using the opposite phase-encoding direction for distortion correction. The off-resonance field was then estimated in FSL (Smith et al., 2004) using a method similar to that described by Anderson et. al. (2003).

### Macaque fMRI acquisition

Macaques were scanned at the Neurophysiology Imaging Facility at the NIH in a 3T Siemens Magnetom Prisma. Whole-brain high-resolution anatomical scans were collected in a single session, under anesthesia, using a single loop receive coil. Several T1 scans were collected and averaged (0.5 mm isotropic resolution, TR = 2.55 s, TE = 2.22 ms, TI = 900 s, FA = 9°), and one T2 was collected (0.5 mm isotropic resolution, TR = 3.2 s, TE = 562 ms, FA = 120°). All functional EPI data were collected using custom-made 8-channel coils (Magnetic Resonance Physics and Instrumentation Group at the Athinoula A. Martinos Center, Massachusetts General Hospital) - on the whole-brain (40-42 slices) - at a voxel size of 1.2 mm isotropic, a TE of 18 ms, a FA of 79°, a partial Fourier of 7/8 and iPAT 2. Repetition times differed between retinotopy (TR = 2 s) and all other experiments (TR = 2.5 s). Most runs were collected in the Head-to-Foot (HF) phase encoding direction (corresponding to AP in the animal’s frame of reference), but we alternated HF and FH for some sessions. In most sessions, we also collected a T1, an in-plane T2, and 10 TRs of spin-echo EPI in the run phase encoding direction and its opposite (HF and FH), to be used for distortion correction (Schallmo et al., 2021).

### Preprocessing

#### Human preprocessing

Functional images underwent distortion, temporal, and motion correction. Opposite phase-encoding acquisitions were processed with FSL TOPUP to estimate susceptibility-induced off-resonance fields, and the resulting correction was applied to the functional images. Distortion-corrected volumes then underwent FSL slice-timing correction using BIDS-derived slice timings and six-parameter FSL MCFLIRT motion correction using mutual-information cost. Framewise displacement was calculated from FSL motion estimates using Nipype’s (Gorgolewski et al., 2011) Power et al. (2012) implementation, and motion estimates were visually inspected.

Motion-corrected runs were registered to native space anatomical images using ANTs (Avants et al., 2009) by rigidly aligning the run-mean EPI to the participant’s native-space T1. These anatomical images were then normalized to the 2009c asymmetric MNI template space using affine and SyN transforms, then the composite transforms were applied to functional data with B-spline interpolation in down sampled anatomical and template grids matched to functional resolution. SPM-derived WM and CSF maps were eroded (3 and 1 iterations, respectively), then combined to define a mask for aCompCor (Behzadi et al., 2007). Five aCompCor components were estimated with Nipype CompCor using polynomial pre-filtering (degree 2). Nuisance models additionally included six MCFLIRT motion parameters and framewise displacement; confounds were z-scored and regressed from coregistered BOLD data.

### Macaque preprocessing

Functional images underwent distortion, temporal, and motion correction. Opposite phase-encoding spin-echo field maps (FH/HF) were paired, affine-resampled when needed, and processed with FSL TOPUP using the macaque-specific b02b0_macaque.cnf configuration to estimate susceptibility-induced off-resonance fields. The resulting correction was applied to functional images. Data then underwent sphinx reorientation, FSL slice-timing correction using BIDS-derived slice timings, and six-parameter FSL MCFLIRT motion correction using mutual-information cost. Frame-wise image quality was assessed by ANTs mutual-information similarity to the run mean, alongside motion estimates and framewise-displacement traces; these measures were summarized in run-level QC reports and videos.

Motion-corrected runs were registered with native space anatomical images using the session mean EPI and session in-plane anatomical image to a subject-level high-resolution anatomical image. A precomputed subject-anatomical-to-NMT v2.1 asymmetric transform was then applied to generate template-aligned functional data at functional resolution.

### Analysis

We introduce Multiview temporal alignment (MVTA) to temporally align the observed macaque MION to the observed human BOLD response during naturalistic stimulation. When combined with hyperalignment (Guntupalli et al., 2018), this Spatiotemporal (ST) Hyperalignment pipeline enables cross-species alignment of representational spaces while accommodating differences in functional topography and hemodynamic response properties.

### Multiview Temporal Alignment (MVTA)

Whereas human BOLD data were acquired using endogenous contrast, macaque data were acquired using the exogenous contrast agent mono-crystalline iron-oxide nanoparticles (MION) to increase the signal-to-noise ratio (Leite et al., 2002). Because the canonical hemodynamic response functions (HRFs) differ between BOLD and MION (Fig. 1D), they can obscure shared representational structure and, if unaccounted for, would introduce a systematic temporal bias: delayed neural responses measured with the faster MION signal could spuriously appear more like earlier responses measured with the slower BOLD signal. For this reason, previous studies (Mantini et al., 2012b, 2012a) correlating responses across species convolved the observed data in each species with the canonical HRF of the other. Although this approach is mathematically valid, the additional convolutional step acts as a temporal filter, discarding high frequency information shared between BOLD and MION (Fig. 1E). This double convolution approach additionally has the potential to temporally-misalign the data if either canonical HRF inaccurately represents the true hemodynamic response, which is known to vary across brain regions (Badillo et al., 2013; Birn et al., 2001; Handwerker et al., 2004, 2012; Lindquist et al., 2009).

MVTA learns a direct transform from MION to BOLD through a multiview alignment framework that maximizes shared multivariate structure across species while deemphasizing species- and individual-specific variance that is unrelated to the transformation of interest and would otherwise bias its estimation. The transformation is initialized from the canonical HRFs (Fig. 1F) and refined through nested cross-validation to minimize discrepancy between cross-species representational spaces (see Supplementary: Hemodynamic Transform Algorithm). The optimized kernel from each training fold was then applied to its corresponding held-out MION data, generating the BOLD-aligned macaque responses used for subsequent hyperalignment analyses. Reported SED is always computed on this held-out stimulus: both the MVTA kernel and the subsequent hyperalignment maps (*w_i_*) are learned from the training fold and are never refit on the test data.

An alternative data-driven approach, temporal kernel CCA (Bießmann et al., 2010), learns an unconstrained spatiotemporal filter, effectively a finite impulse response (FIR)-like model in which separate weights are estimated for multiple temporal lags and voxels. While flexible, this approach heavily increases model complexity and can risk overfitting. In contrast, ST-hyperalignment constrains spatial transformations to be strictly time-invariant, modeling temporal filtering as a physiological consequence of the hemodynamic response (i.e., the HRF) acting as a slow, blood-based proxy for neural activity. By enforcing time-invariant spatial transformations, ST-hyperalignment reduces free parameters to stabilize estimation while preserving a physiologically meaningful separation between the spatial organization of population responses and their hemodynamic temporal properties.

To date, hyperalignment (i.e., multiview analysis applied to neuroimaging data) has been used primarily to identify representational structure shared across homogeneous groups of subjects (Haxby et al., 2011, 2020). Here we use the Shared Response Model (SRM; Chen et al., 2015) implemented in the nltools python package (Computational Social Affective Neuroscience Laboratory, 2015/2026). The SRM approximates each subject’s data, *X_i_*, as the product of a shared representational space, *S*, and a subject-specific projection matrix, *w_i_*, together with a residual term, *E_i_*:

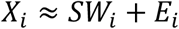

Importantly, *E_i_* includes not only noise, but also signal unique to individual subjects. The SRM is statistically powerful because it identifies stimulus-driven structure that is shared across subjects, while treating signals that do not align as residual. After fitting, subject *i*’s coordinates in the shared space are the projection:

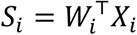

The shared response estimated during training is the average of these projections 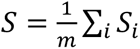. We fit *w_i_*_(*tr*)_ on training responses and then transformed each subject’s held-out test data into the shared space, 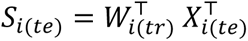 without refitting *w_i_*.

Human and macaque training data were jointly used to estimate a latent common space, yielding a subject-specific weight (or projection) matrix for each participant (Fig. 1B). These learned projection matrices were then applied to each subject’s held-out test data, translating subject-specific voxel data into the axes of the latent common space.

Human and macaque training data were jointly used to estimate a latent common space, yielding a subject-specific weight (or projection) matrix for each participant (Fig. 1B). These learned projection matrices were then applied to each subject’s held-out test data, translating subject-specific voxel data into the axes of the latent common space.

### Training and testing

Both stages were trained and tested on independent data using cross-validation across three movies and two static image sequences. By default, models were trained on four stimuli and tested on the fifth. When directly comparing movies and static image sequences, we used balanced-stimulus training to mitigate training bias, fitting each method on one movie and one static image sequence before testing on each held-out stimulus in turn. For between-species analyses, between-subject cross-validation yielded 12 independent subject-level SED estimates (nine human, three macaque) for group inference. Within each training fold, hyperalignment was fit to species-balanced subsets (three humans and three macaques), with humans bootstrap-resampled to match the macaque sample size and SED averaged across macaque repetitions. This balancing prevented either species from disproportionately influencing estimation of the shared space.

### Stimulus epoch decoding

We quantified shared information using within- and cross-species stimulus epoch decoding (SED; Borovska & De Haas, 2024; Busch et al., 2021; Haxby et al., 2011b; Van Uden et al., 2018). For a held-out subject, temporally corresponding epochs should correlate more strongly with the comparison-group average than noncorresponding epochs. We therefore correlated all nonoverlapping 10-TR epochs and classified each epoch by its maximally correlated counterpart. Accuracy was averaged across row- and column-wise classification to produce a symmetric measure. Chance is the reciprocal of the number of nonoverlapping epochs in the tested stimulus. Within-species SED compared each subject’s projected test data with the mean projected test representation of all other subjects from the same species. Between-species SED compared each subject’s projected test data with the mean projected test representation of all subjects from the other species. Each subject therefore contributed a view of the latent common representational space for each held-out stimulus response.

Group-level effects were first assessed using one-sample *t*-tests across subject-specific SED values. Given the modest sample size, we additionally report non-parametric Wilcoxon signed-rank tests, which make fewer distributional assumptions and provide a robustness check on the parametric t-tests. For dense ROI-by-ROI and cortical-map analyses, displaying every result in the figure was impractical. Instead, we used a highly conservative reporting criterion, requiring both that every subject’s SED exceeded chance and that the mean SED across subjects was significantly greater than chance by a binomial test. This criterion is analogous to combining a one-sided sign test, requiring all subjects to show above-chance SED, with a binomial test of the group-mean decoding rate. It therefore emphasizes effects that are both reliable in direction across subjects and statistically above chance at the aggregate level. Additional results are provided on the accompanying website.

### Quantifying organization among V1–V4

The V1–V4 results were summarized with a subject-level ordinary least-squares model fit separately to the within-human, within-macaque, and cross-species shared-information matrices. Each matrix was hemisphere-resolved (left and right V1, V2, V3, and V4; 8×8). Cross-species matrices had macaque areas as rows and human areas as columns. Each cell was the shared information for that area pair, averaged across runs of one stimulus type. The static-image matrix and the movie matrix were stacked, and shared information was z-scored within subject across that stack. The primary models used the default hyperalignment training (four of five stimuli). The within-species predictors were representational fingerprint, hierarchical proximity, and hemisphere match, together with stimulus condition and its interaction with each of those predictors. The cross-species model added a triangle-bias term and its interaction with condition. The same cross-species model was refit after excluding V4.

Fingerprint contrasted same-name with different-name area pairs, regardless of hemisphere. Hierarchical proximity, defined only for different-name pairs, was the negative absolute difference in assigned hierarchy level, such that larger values indicate closer positions in the hierarchy. Hemisphere match contrasted same-with opposite-hemisphere pairs. Triangle bias, used only cross-species, was defined with macaque areas as rows and human areas as columns. Pairs in which the macaque area was higher order than the human area formed the positive side, and the swapped pairs formed the negative side. Same-name cells were omitted from hierarchical proximity and from triangle bias. For each binary contrast, the positive side was coded +1 and the negative side was coded in proportion to the number of positive cells, so that the contrast summed to zero. Every predictor except condition was then z-scored on the cells where it was defined; omitted cells were set to the mean of that standardized predictor. Coefficients on these predictors are therefore associations in within-subject standardized units, not raw differences in shared information and not changes per hierarchy step.

Condition was coded −0.6 for static images and +0.4 for movies. These weights match the two static image sequences and three movies, and they place the fingerprint, hierarchy, hemisphere, and triangle coefficients on that weighted combination of the two stimulus types. Because the two codes differ by 1, each condition interaction equals the movie-minus-static change in that effect. Again, condition itself was not z-scored.

Human within-species inference used one-sample t-tests and Wilcoxon signed-rank tests on the subject-level coefficients. Macaque coefficients were summarized by their range across the three animals. Cross-species inference used the same two tests on the combined set of human and macaque subject-level coefficients. Each subject contributed one cross-species matrix; the test was not performed across human–macaque pairs.

As a separate summary of hemispheric specificity, each subject’s within-species 8×8 matrix was scaled to sum to 1 and reduced to the 10 unique unordered area pairs (including same-name pairs). For each pair, the effect was (matched − mismatched) / (matched + mismatched), and these ratios were averaged. Humans and macaques were compared with a one-tailed Welch test and a species-label permutation test. The same comparison was also run separately for each of the 10 pairs, with false discovery rate correction across pairs. Finally, the cross-species regression was repeated on shared-information matrices from a control in which hyperalignment was trained on one movie and one static-image sequence.

For the Fig. 4B inset, we computed a matched-hemisphere triangle-bias summary on shared-information (SED) matrices from hyperalignment trained on one movie and one static-image sequence in each fold (balanced training). Movies and static images were analyzed separately. For each subject and stimulus type, cross-species cells were averaged into a 4×4 matrix (macaque rows, human columns), mean upper-triangle and lower-triangle values were taken, and the subject-level index was (upper − lower)/(upper + lower). Group inference used one-sample t-tests of each condition against zero and paired t-tests comparing movies with static images. The inset shows V1–V4.

### Joint multidimensional scaling

In the final analysis of V1–V4 (Fig. 4C), we performed two-dimensional multidimensional scaling (Borg & Groenen, 2005) to summarize the pattern of inter-areal relationships in a joint within- and cross-species space. Distances were calculated from a combined within- and cross-species distance matrix (distance = 1 − SED, averaged across subjects).

### Eye movements and neural similarity

Differences in scanner eye-tracking configuration impacted data yield. Whereas macaque eye tracking was performed with a direct line of sight from outside the bore, human tracking required participants to view the stimulus through a mirror within the head coil. Because this configuration is more susceptible to pupil-tracking loss, blink artifacts, and calibration drift, some human data was excluded after quality screening (macaque data remained complete; each human contributed data for at least one movie). This yielded a final dataset of 24 cross-species subject pairs for Forrest, 18 pairs for Growth, and 21 pairs for the macaque movie. Eye-movement similarity was calculated as the average between-subject Pearson correlation of horizontal and vertical gaze time series, and neural similarity was defined as the Spearman correlation of subjects’ responses within a shared 50-dimensional Shared Response Model space (SRM; Chen et al., 2015). We assessed the relationship within each visual region of interest (V1–V4) separately using ordinary least squares regression of neural similarity on eye-movement similarity. A single slope was fit across subject-movie pairings, so differences among movies remained in the slope. Reported coefficients were obtained by applying a Fisher-z transform to eye-movement and neural similarity with each ROI prior to model fitting.

Because subject–movie pairings were not independent, with each subject contributing observations for multiple movies, significance was assessed using a permutation test that preserved this dependence structure. Because movie labels could be exchanged only for human– macaque pairs with eye-tracking data for all three movies, both the permutation test and the corresponding unpermuted model were restricted to this subset. Neural similarity remained fixed to its original subject-movie pairing, whereas eye-movement similarity was reassigned according to the permuted human and movie labels while macaque identity was held fixed. All 4,320 possible permutations were enumerated and applied identically across V1—V4. In every region, the observed slope fell in the upper tail of the permutation distribution (one-sided permutation *q* < 0.05).

### Atlas-wide correspondence

We used the Glasser HCP atlas for human (Glasser et al., 2016) and the CHARM (level 4) atlas for macaque (Jung et al., 2021). We learned a distinct HRF transformation for each between-species atlas pair. For an ROI pair to be considered significant, the binomial *p*-value associated with the mean SED across subjects had to survive FDR correction (Benjamini & Hochberg, 1995; *q* < 0.05), and the cross-species SED had to exceed chance for every individual subject. The matrix in Fig. 6 was sorted so that the strongest SED values were shifted towards the top left corner, then divided by lobe and hemisphere. For the cortical surfaces in Fig. 7A, the color of each macaque ROI is the maximum SED value across the corresponding row of that matrix, and the color of each human ROI is the maximum SED value across the corresponding column.

### Cortical expansion

We performed a spin test (1000 permutations) using the Neuromaps python package (Markello et al., 2022) to assess global spatial correlation between our SED-derived estimate of information sharing (sign-reversed cross-species SED values of the unthresholded map) and the independent surface expansion map (J. Hill et al., 2010). Each location’s signed contribution to the global correlation was the element-wise product of the z-scored surface maps.

### Cross-species searchlights

Macaque searchlights were implemented on the cortical surface to preserve the intrinsic geometry of the cortical sheet (geodesic distances). We used a 6mm searchlight radius. Human seed regions were defined volumetrically to minimize spatial resampling. To minimize computational demands, we re-used the atlas-derived transforms from the atlas-based analyses. To calculate the HRF transform for each cross-species pair, we first identified the atlas regions represented within the human seed, then within each macaque searchlight sphere, and computed a weighted average of those transforms, with weights proportional to the number of vertices assigned to each region. The right temporoparietal junction seed was the FDR-corrected (*q* < 0.01) Neurosynth association test map for the term “mind” (Yarkoni et al., 2011). The word seed used individual-specific functional masks from the human word localizer (words and consonant strings > checkerboards).^4^ Fusiform face area seeds were defined for each subject using the category localizer (faces>objects). Parahippocampal place area seeds were also defined for individual subjects using the category localizer (places> objects). The frontal eye field seed was defined using the Glasser atlas (2016).

## SUPPLEMENTARY

### Supplementary Discussion

#### Multiview Temporal Alignment (MVTA)

MVTA seeks to learn a convolutional kernel that aligns MION data directly to BOLD. We compared it to several alternatives using simulated data with known ground-truth structure. A central challenge is that observed fMRI signals potentially reflect mixtures of multiple sources of variance (e.g., components shared across species, components shared only within a species, and components unique to individual subjects). Only the shared component is informative for learning cross-species temporal transformations, but unshared components may obscure the shared structure needed to learn a meaningful transform.

We generated five synthetic fMRI runs for two “macaque-like” and two “human-like” datasets, each comprising multivoxel responses driven by multiple latent neural signals.

First, we generated latent neural time series for 250 synthetic voxels from three classes of signals: (i) two signals shared across species (“shared”), (ii) two signals shared only within species (“group-specific”), and (iii) two subject-specific signals. Neural signals (*X_neural_* ∈ *R^T^*^×*N*^*^signals^*) were generated as random binary sequences (Bernoulli process). In real data, the relative contributions of these signal classes are unknown. Our goal was to include sufficient unshared neural variance to mimic the species- and individual-specific representational structure that we expect in many regions of the brain.

Second, we then combined these signals using a random weighted average, where weights were sampled from a uniform distribution and normalized to sum to one. These voxel weights were stable across runs within a subject but differed between subjects. We next convolved the neural time series of each voxel with the species-specific canonical HRF (MION or BOLD). The resulting voxel time series were then z-scored.

Finally, we compared four different methods to account for the temporal misalignment introduced by the different HRFs:

1. Double convolution. Following Mantini et al. (2012b, 2012a) we convolve each species’ data with the other species’ canonical HRF, placing both time series in an intermediate’ temporal space. This approach enables voxel-wise comparisons but imposes strong low-pass filtering. Reliance on canonical HRFs is also subject to compounded HRF mis-modeling error, which we do not consider here.
2. Multiview without temporal alignment. We fit a variance-partitioning multiview model (the Shared Response Model; Chen et al., 2015) directly to untransformed BOLD and MION data, projecting view-specific data into a shared latent space. This provides a baseline test of the degree to which multiview representational alignment can recover cross-species structure without correcting the temporal mismatch between MION and BOLD.
3. Temporal alignment without multiview. We learned a convolutional kernel that maps MION-to-BOLD by minimizing a voxel-space mismatch objective (also sum of squared errors) between MION and BOLD voxel timeseries. This approach tests whether a data-driven temporal transform can be recovered without partitioning unshared variance.
4. MVTA.

Together, these methods form an ablation grid designed to isolate the core methodological factors of interest: canonical versus learned temporal alignment, and the absence versus presence of multiview variance partitioning.

| Method | Temporal Alignment | Variance Partitioning |
| --- | --- | --- |
| 1. Double convolution | Canonical HRFs | No |
| 2. Multiview without temporal alignment | None | Yes |
| 3. Temporal alignment without multiview | Learned | No |
| 4. MVTA | Learned | Yes |

We evaluated these methods in two ways. First, we evaluated temporal alignment on 10,000 held-out neural time series. For each, we compared the BOLD-convolved signal with the corresponding signal produced by each alignment procedure; for the no-alignment condition, this was the MION-convolved signal. Performance was measured by correlation with BOLD (Supplementary Figure 1A). Second, to estimate shared stimulus information, we performed Stimulus Epoch Decoding (SED) in the SRM-derived common space representation of the held out stimulus (Supplementary Figure 1B).

The two analyses yielded similar results—MVTA-corrected MION timeseries performed best, strongly outperforming alternatives. Applying the multiview approach without any temporal alignment performed reasonably well (as the SRM can accommodate the difference in the sign of the HRFs; Figure 1D). We caution against using this approach, however, as differences in the timing of the peak HRF signal would introduce systematic temporal bias. Delayed neural responses measured with the earlier-peaking MION signal would spuriously appear more like earlier responses measured with the slower-peaking BOLD signal.

**Supplementary Figure 1.**
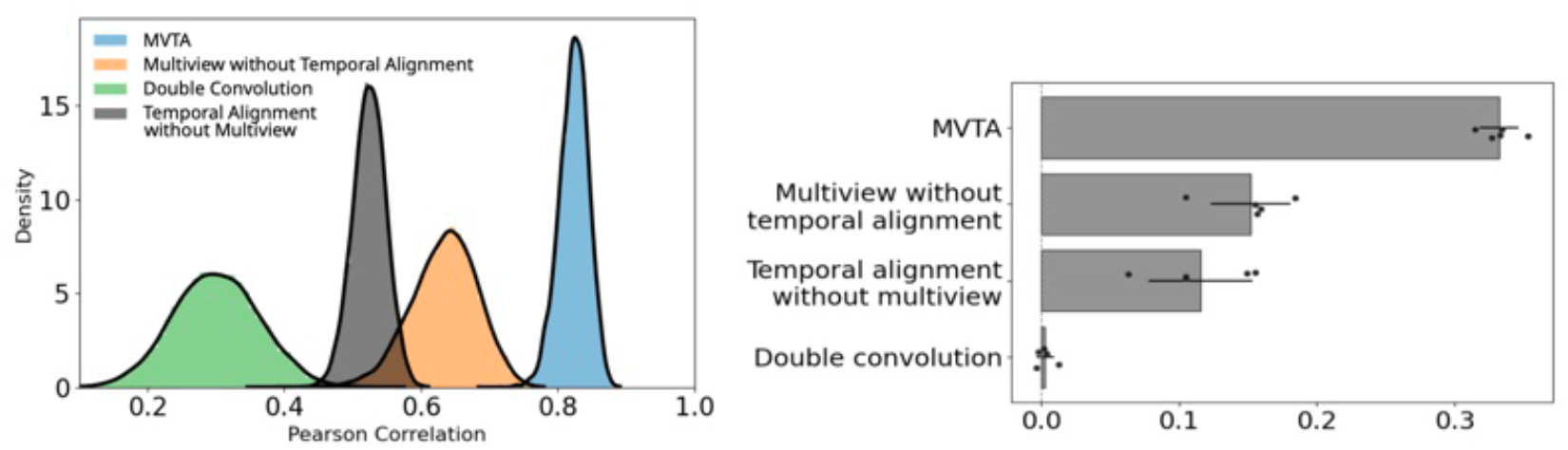
**A)**: Distribution of correlations between ground-truth BOLD and transformed MION signals across 10,000 held-out simulated neural time courses for each learned HRF method. MVTA (blue, mean r=0.82) substantially outperformed multiview without temporal alignment (orange, mean r=0.63), temporal alignment without multiview (black, mean r=0.52), and double convolution (green, mean r=0.3).Black lines depict kernel density estimates of the underlying histograms. **B)**: Shared stimulus information (SED) between BOLD and transformed MION in shared space. Mean accuracy across folds (error bars: +- 1SD; points: individual folds). MVTA substantially outperformed alternative approaches.

### Hemodynamic Transform Algorithm (MVTA)

An HRF transform was learned for each between-species ROI pair through a multiview approach using multivoxel movie and temporally synchronized x static image data. To promote generalization and prevent overfitting, the transform was trained and evaluated using nested cross-validation. We used a fixed shared space dimensionality of 50. In principle, individual subject transforms could be learned, but we opted to learn a single transform for each ROI pair to reduce computational cost while accounting for differences between species and brain-regions. Learning a transform from BOLD to MION would be more complicated due to its later signal onset. Code demonstrating the algorithm on simulated movie responses is available at https://github.com/kbraunlich/multiview_temporal_alignment.

Because the MION response typically precedes the onset of the BOLD response, learning the reverse transformation (BOLD to MION) is fundamentally more difficult as it requires predicting the MION signal that occurs prior to BOLD onset. In cases where it is unknown which modality’s temporal filter responds first, the mapping can simply be fit in both directions on training data, allowing the framework to empirically select the path that yields the best temporal alignment.

1. **Outer loop cross-validation**: Divide the training set stimuli into *K* folds. For each stimulus *kᵢ* in the *K* folds:

**a.** Assign stimulus *kᵢ* to the outer fold test set, and the others to the outer fold training set.
**b.** Create an informed guess for the transformation from MION to BOLD^5^

i. Estimate a fixed SRM dimensionality using the outer folder training data only. Transform MION data using the canonical MION-to-BOLD transform, fit a probe SRM. Optionally compute the fractional temporal energy of each shared component and retain components with at least the uniform expected share of energy (1/*K;* at least one component is selected).
**c. Inner loop cross-validation**: Divide the outer-fold training set stimuli into *K–1* folds. For each *kⱼ* in the *K–1* folds:

i. Convolve outer loop training set MION data with the proposed transform.
ii. Assign stimulus *kⱼ* to the *inner loop* test set, and the others to the *inner loop* training set:

– Fit multiview model using the fixed SRM dimensionality on *inner loop* training set.
– Transform *inner loop* test set data to shared space.
– Calculate the average sum of squared errors between the BOLD and MION *inner loop* test set shared spaces.
**d.** Repeat Step 1c, each time modifying the proposed transformation to minimize the average between-group disparity, until convergence (using BFGS).
**e.** Apply the learned transformation to outer-loop training and test MION (kernel estimated without the test stimulus).
**f.** Hyperalignment is fit on the outer-loop training stimuli and then used to transform the held-out test stimulus into the shared space; SED is computed on this test-set projection.
2. Calculate average movie segment decoding accuracy across *K* outer loop folds.

### Temporal Differences across V1-V4 and MT

In Supplementary Figure 2, we plot the MVTA-derived transforms between macaque V1, and human V1—V4 + MT. Relative to the transforms involving macaque V1-V4 (which are highly similar), the transform involving macaque MT is delayed in time. Although this is consistent with MT’s known role in motion processing, this possibility cannot be distinguished from a regional difference in hemodynamic response shape.

**Supplementary Figure 2.**
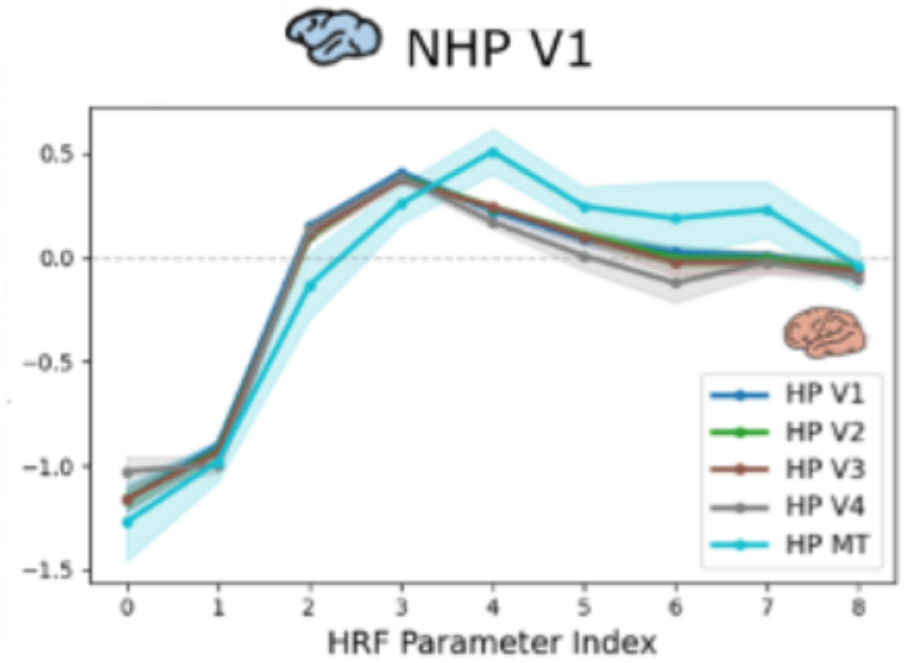
Learned HRF transform between pRF-defined macaque V1 and human V1, V2, V3, V4, and MT (TR = 2.5 s). Shaded regions indicate ±1 SD across runs (three movies and two static image sequences). The learned transform was largely consistent from macaque V1 to human V1–V4, whereas the transform to human MT showed a more gradual rise and a later peak. It is unclear whether this difference reffects neural timing or regional variation in the HRF.

### Cross-Species Dorsal and Ventral Components of V1-V4

Retinotopic visual areas V1, V2, and V3 are organized into separate representations of the upper and lower visual fields (the extent to which this is also true for V4 is debated; Goddard et al., 2011; Hansen et al., 2007; Wade et al., 2008; Winawer et al., 2010). Here we ask: Does cross-species correspondence preserves the upper-lower visual field representational structure? We split each pRF-derived ROIs into their dorsal and ventral components and repeated the between-species analyses (Supplementary Figure 3). After calculating shared stimulus information (SED) values for all ROI-pairs, we fit regression models to each subject. Group-level inference was then performed on the resulting subject-wise regression coefficients using one-sample *t*-tests and Wilcoxon signed-rank tests. As with the full pRF-derived ROIs, we included terms for stimulus (movies vs. static images), hierarchical bias (upper vs. lower heatmap triangle), representational fingerprints (diagonal vs. off-diagonal), and canonical hierarchical structure. However, we additionally included a regressor for dorsal/ventral asymmetry (matched vs. mismatched dorsal ventral components). Finally, we included all two-way interactions with stimulus type. Although we include human V4 in the heatmaps of Supplementary Figure 3, we exclude it from statistical analyses due to uncertainty about the relative organization of its dorsal component between species (Goddard et al., 2011; Hansen et al., 2007; Wade et al., 2008; Winawer et al., 2010).

**Supplementary Figure 3.**
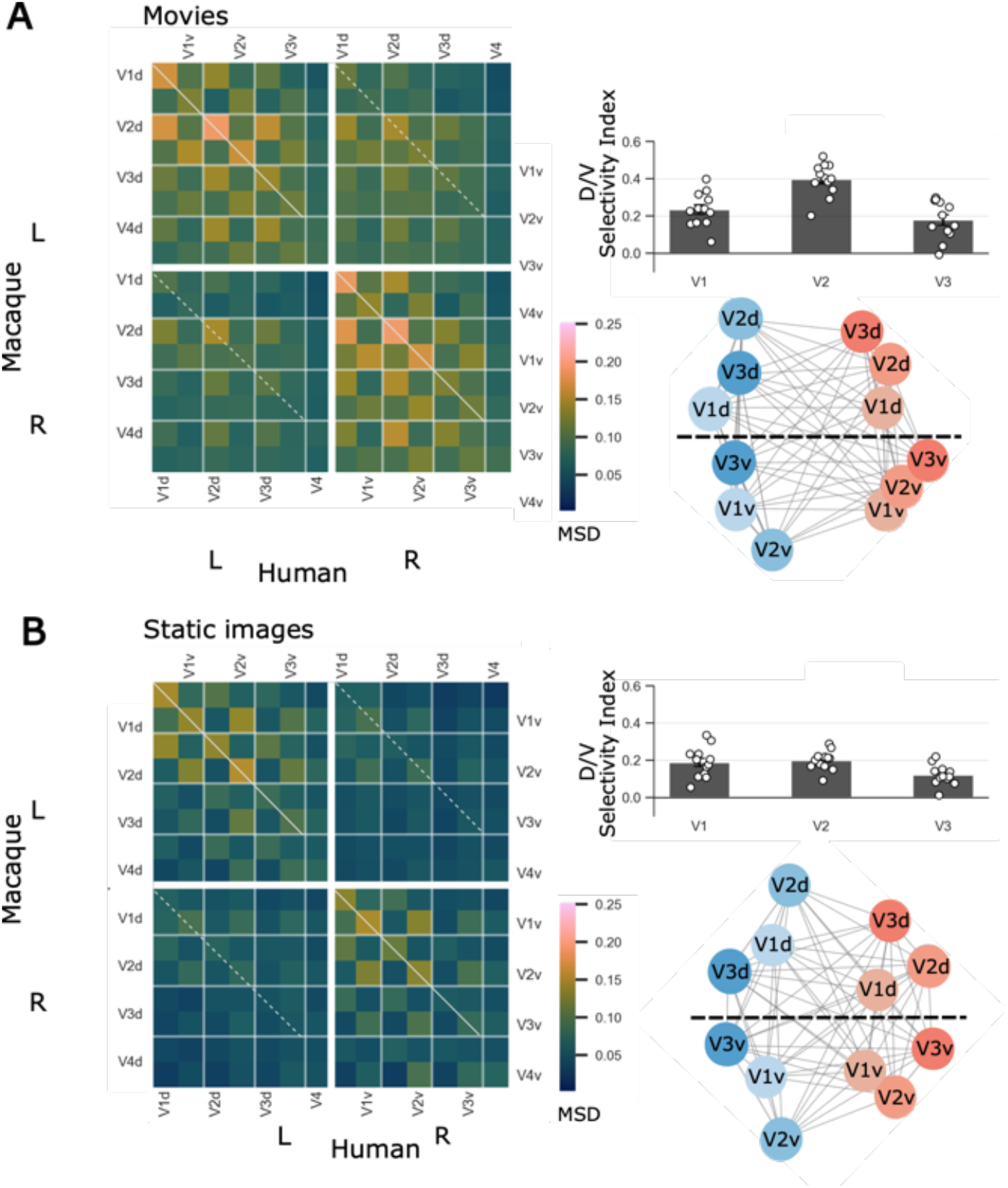
**(A)** Movie viewing (left): Heatmaps of cross-species SED during naturalistic movie viewing. Representational similarity is stronger between matched dorsal and ventral subdivisions. The matrices display the hierarchical asymmetry effect—where higher-order macaque regions share more information with lower-order human regions (e.g., macaque V3/V4 sharing high similarity with human V1/V2). A hemisphere bias is also present: SED between macaque V3/V4 and human V1–V3 is greater for the left-macaque/right-human pairing than for the reverse. White lines depict the identity diagonal (solid = matched hemispheres; dashed = mismatched hemispheres). Dorsal/ventral selectivity index **(top right):** Mean dorsal/ventral (d/v) selectivity index across participants for visual areas V1, V2, and V3 (V4 is excluded). For each participant and visual area, four SED values were extracted from the movie-viewing cross-species matrix by first averaging over matched hemispheres (left-to-left and right-to-right), then averaging over the dorsal and ventral components within each subdivision category: matched = mean of dorsal-to-dorsal and ventral-to-ventral; mismatched = mean of dorsal-to-ventral and ventral-to-dorsal. The d/v selectivity index was defined as (match − mismatch) / (match + mismatch). An index of zero indicates no subdivision preference; positive values indicate greater SED accuracy for subdivision-matched than subdivision-mismatched region pairs. Bars show the group mean; error bars indicate the standard error of the mean. Individual participant values are overlaid as points**. (B)** Static images (left): Heatmaps of cross-species SED during static image viewing. Under these static stimulus conditions, the hierarchical asymmetry effect is not apparent. Multidimensional **scaling (bottom right):** Joint within- and cross-species MDS space. The two dominant dimensions primarily capture species differences (horizontal axis) and dorsal-ventral representational differences (vertical axis), as these binary classes are linearly separable. Although the hierarchical organization is somewhat obscured in this two-dimensional projection, it remains apparent in the full heatmaps.

As with the full ROI analysis, we found that shared stimulus information (SED) during movies tended to be greater than during static image sequences (*t*(11) = 10.04, *p* <0.001, *W* = 0, *p* <0.001). The main hemispheric effect was not significant (*t*(11) = 0.47, *p* = 0.65 *, W* = 27, *p* = 0.38), however, the hemispheric effect was significantly stronger during movies (*M* β = 0.3, SEM β = 0.05) than during static images (*M* β = 0.01, SEM β = 0.03) the interaction was significant (*t*(11) = 6.11, *p* < 0.001 *, W* = 1, *p* < 0.001). Diagonal values (*M* β = 0.36, SEM β = 0.04) were significantly greater than off-diagonal values (*t*(11) =3.39, *p* < 0.001, *W* = 0, *p* < 0.001), indicating distinct representational fingerprints. This effect did not differ between stimulus types (*t*(11) = - 0.62, *p* = 0.54*, W* = 33, *p* = 0.68). Although this pattern is not readily apparent in the two-dimensional multidimensional scaling projections (Figure 1G), which provide only low-dimensional visualizations of the full distance structure, the off-diagonal values (*M* β = 0.17, SEM β = 0.016, *t*(11) = 10.6, *p* < 0.001 *, W* = 0, *p* < 0.001), and this effect was not significantly different between movies and static image stimuli (*t*(11) = 2.02, *p* = 0.07*, W* = 16, *p* = 0.08). SED tended to be stronger for matched dorsal/ventral components than mismatched components (*M* β = 0.46, SEM β = 0.046, *t*(11) = 9.91, *p* < 0.001*, W* = 0, *p* < 0.001), and this effect did not differ between stimulus types (*t*(11) = 0.34, *p* = 0.7*, W* = 35, *p* = 0.8).

The two MDS axes discriminate species and the dorsal/ventral split.

### Methodological Considerations and Contributions

A practical consideration concerns behavioral training requirements. While we trained the first 2 NHPs (Ri and Mo) to perform a 1-back task during the localizer tasks and the static image stimulus, we subsequently added a 3rd macaque subject (Wo) without the behavioral task. We found that results largely generalized to this macaque, suggesting that (at least for the brain regions identified here) behavioral task is not strictly necessary for reliable localization and identification of shared signal.

**Supplementary Figure 4.**
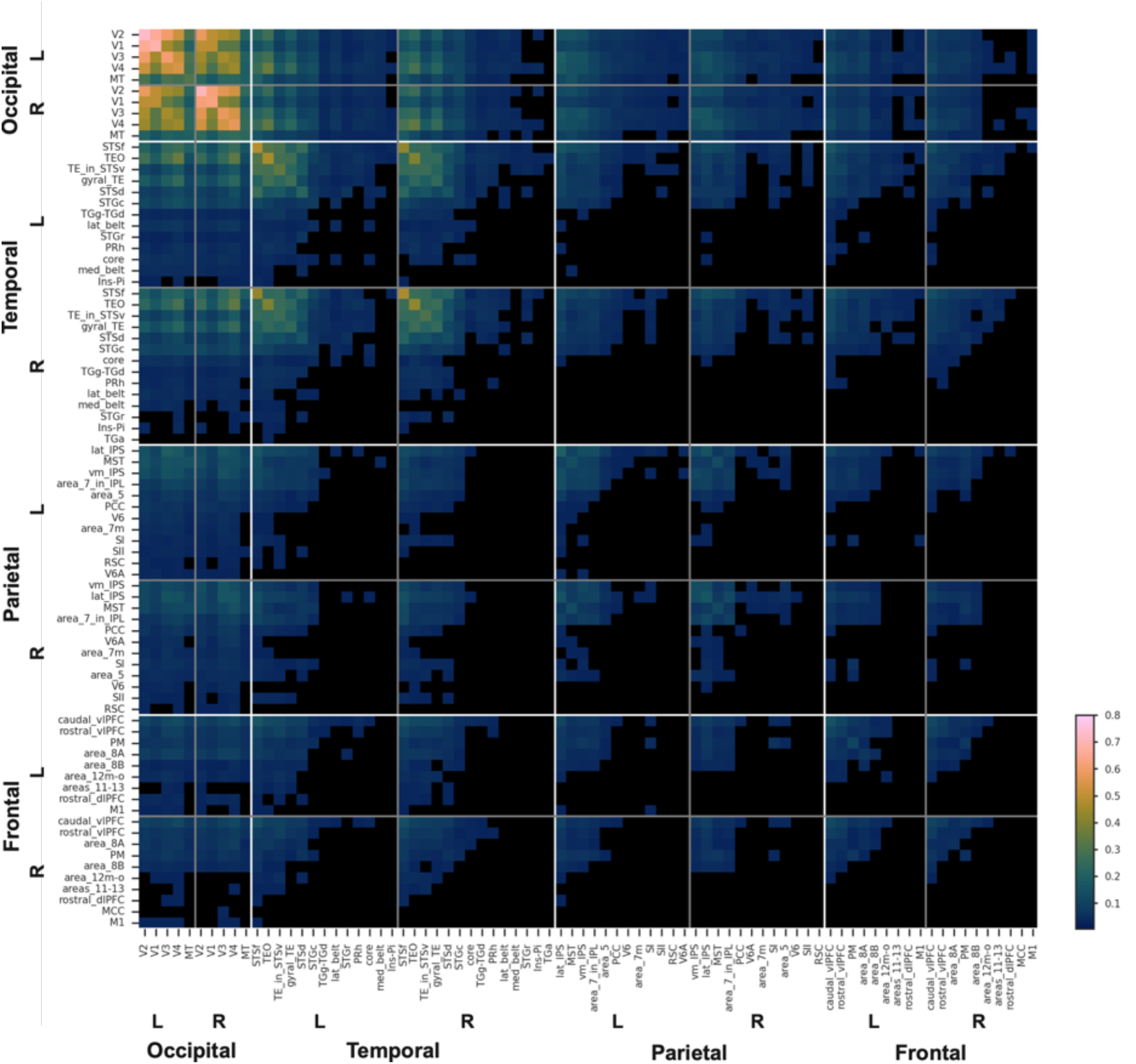
Macaque CHARM (4) atlas average SED accuracy across all stimuli. (FDR corrected mean binomial test q<0.05 and all subject SED >chance.)

**Supplementary Figure 5.**
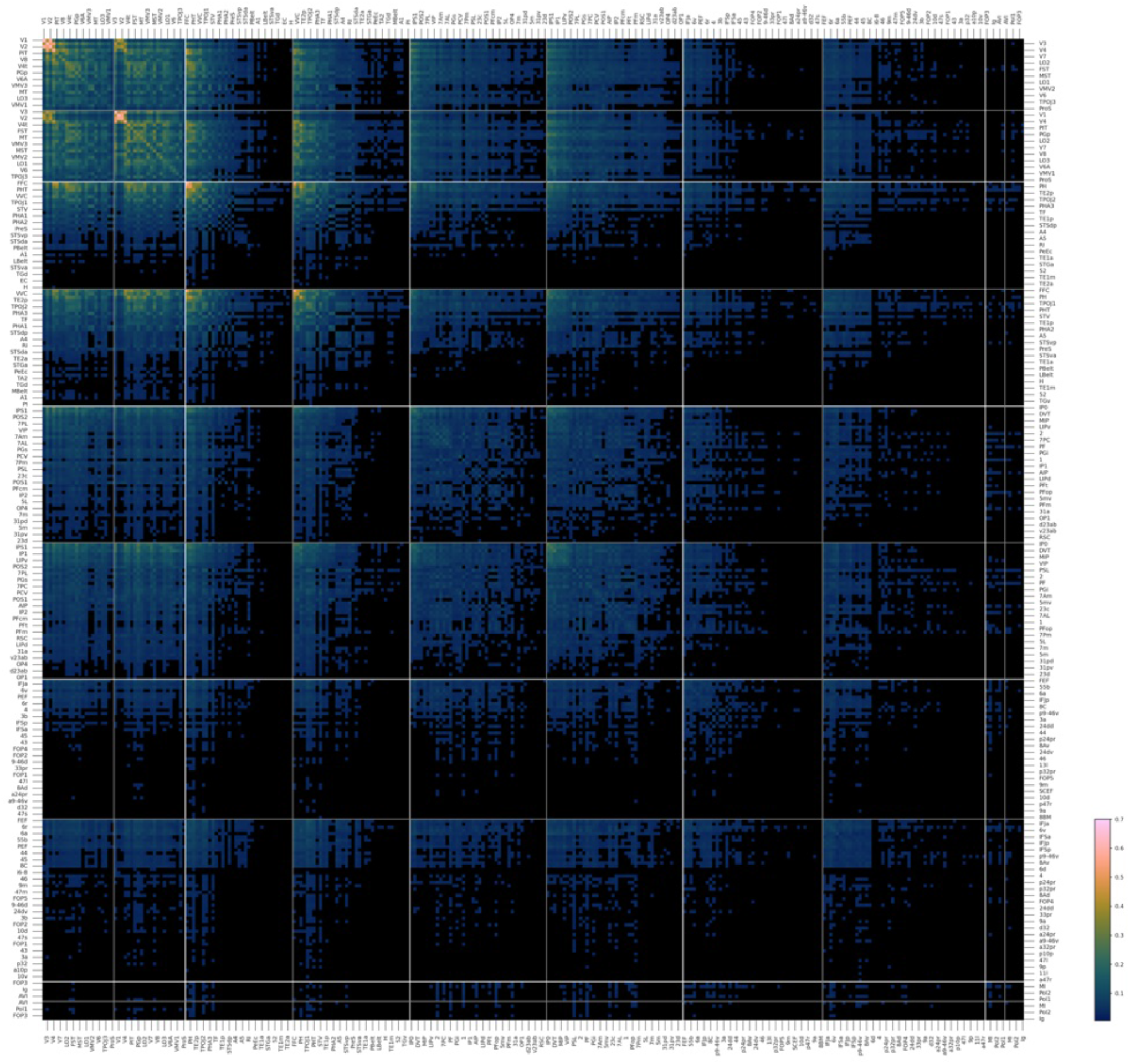
Human Glasser atlas average within-HP SED accuracy across all stimuli. Note that odd and even numbered labels alternate between top/bottom and left/right axes. (FDR corrected mean binomial test q<0.05 and all subjects>chance.)

## Human Brain Coverage

**Supplementary Figure 6.**
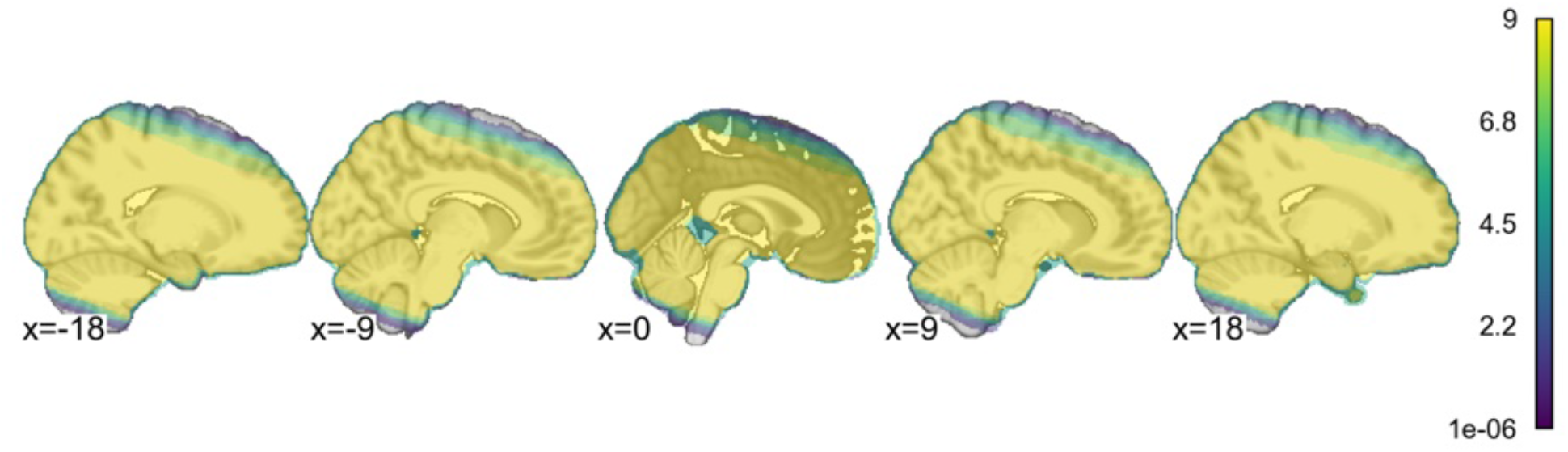
Human brain coverage. Colormap indicates the number of subjects with coverage at each voxel.

**Supplementary Figure 7.**
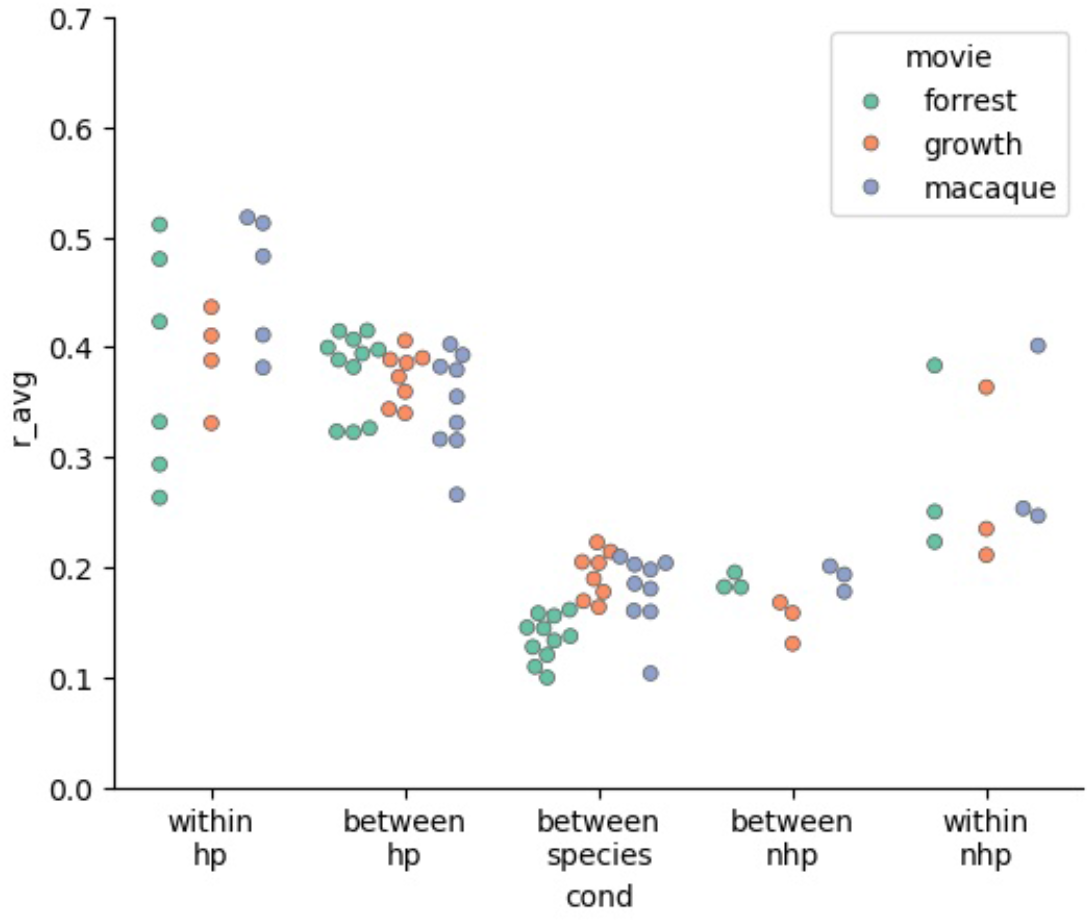
Pearson correlation coefficient between eye-movement traces. Within HP and NHP are reliability within individual subjects. Between HP and NHP are reliability across subjects within each species. Between species is across species. Not all HP subjects had reliable eye-tracking data for all stimuli.

**Supplementary Figure 8.**
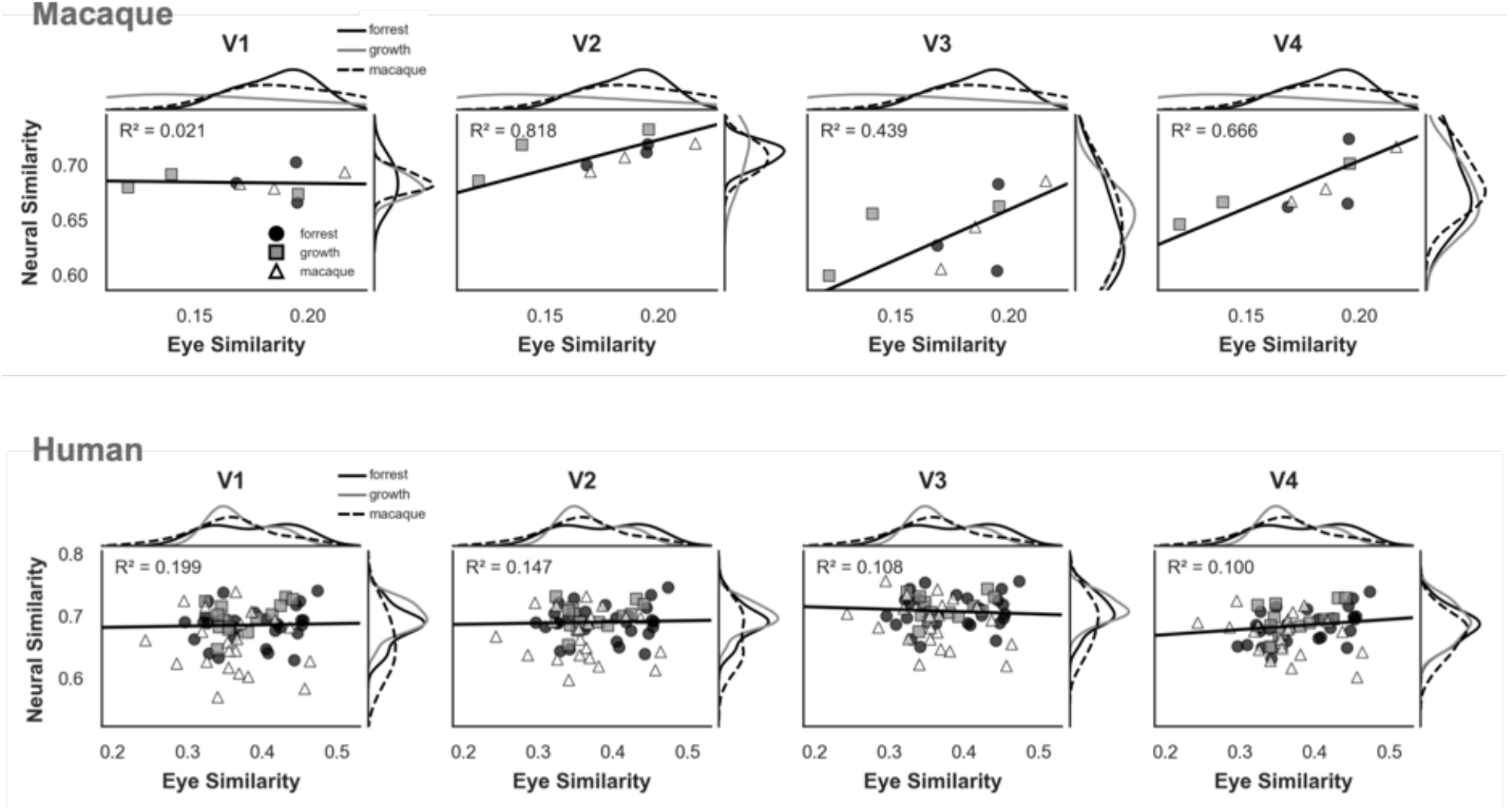
Within-species relationship between eye-movement similarity and neural similarity in early visual cortex. Scatter plots show eye (Pearson correlation) and neural similarity (Spearman correlation) for each human-macaque subject pair. Marginal distributions depict the corresponding similarity values. Filled circles: Forrest Gump; filled squares: Growth; open triangles: Macaque clips. Black lines indicate least-squares regression slopes in the original units (unstandardized). Positive slopes indicate that subject pairs with more similar eye movements exhibit more similar neural responses. Because of three statistical challenges, we treat these analyses as descriptive and do not estimate statistical significance. 1) As each subject contributes to multiple ROI pairs, they are not independent. 2) With N=3, the macaques have an insufficient number of pairs. 3) In humans, the uniformly high similarity of eye-movement traces limits sensitivity to an association with neural similarity.

**Supplementary Table 1:**
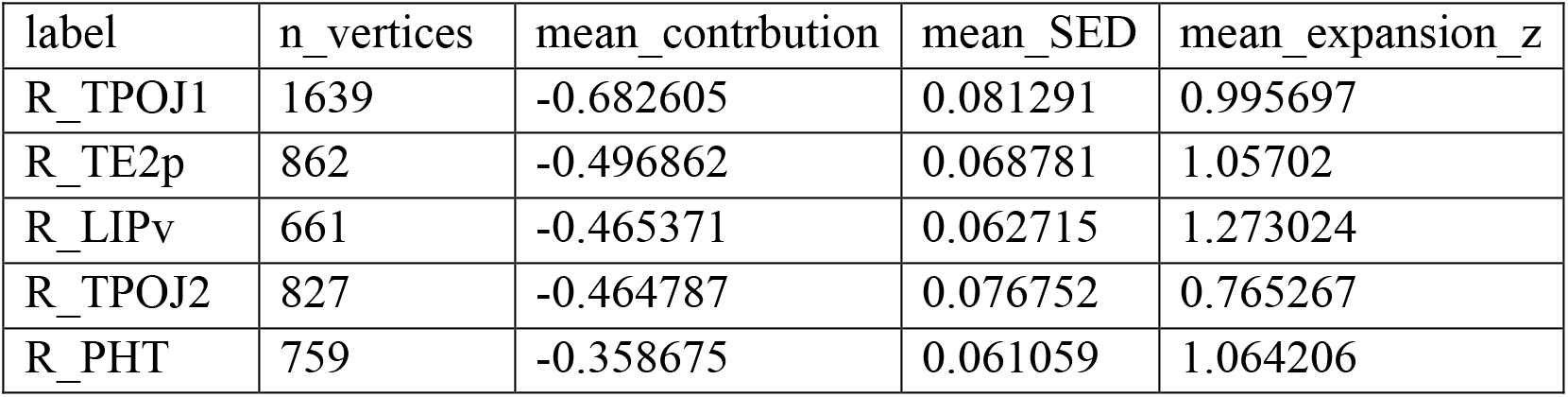
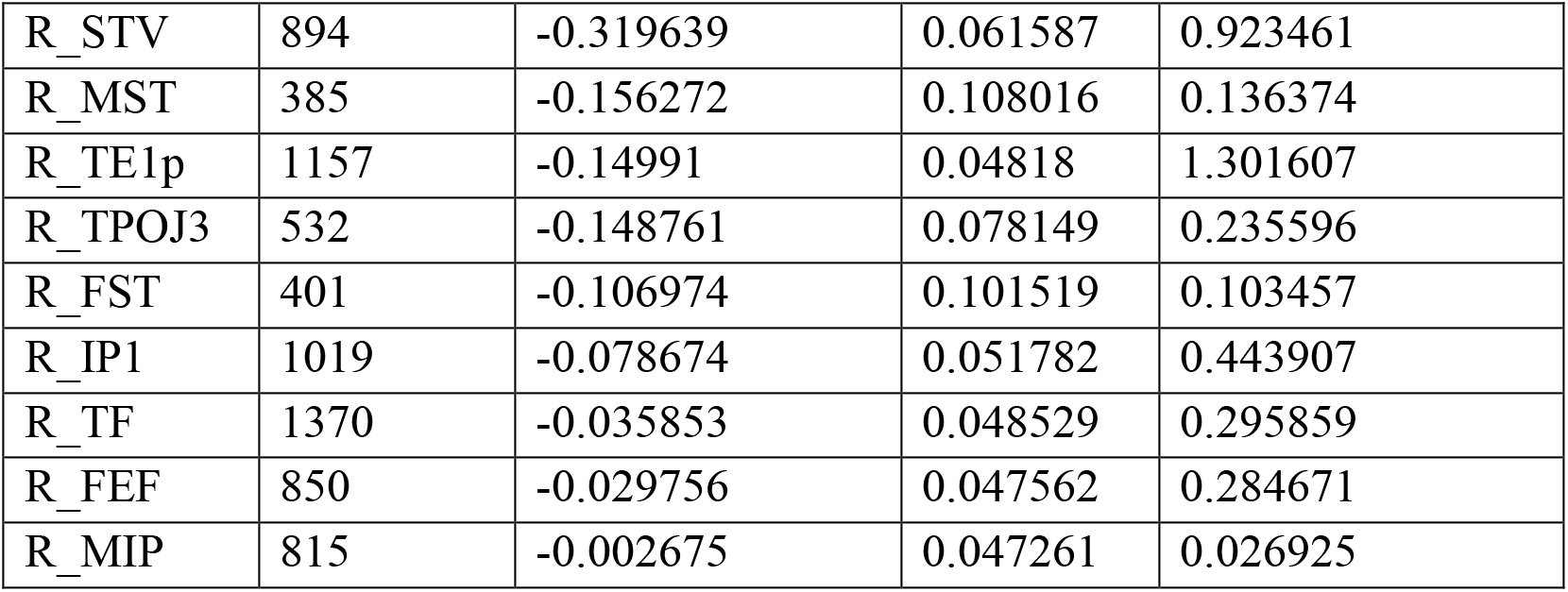
Regions with high expansion and high information sharing (negative contribution to the global correlation). Although regions with high cross-species information sharing tended to localize to areas associated with low cortical expansion, several regions showed high cross-species information sharing despite substantial cortical expansion. These regions may reflect strong selective pressure to preserve function during cortical expansion or, alternatively, an increased allocation of cortical resources to support those functions.

| label | n_vertices | mean_contribution | mean_SED | mean_expansion_z |
| --- | --- | --- | --- | --- |
| R_TPOJ1 | 1639 | -0.682605 | 0.081291 | 0.995697 |
| R_TE2p | 862 | -0.496862 | 0.068781 | 1.05702 |
| R_LIPv | 661 | -0.465371 | 0.062715 | 1.273024 |
| R_TPOJ2 | 827 | -0.464787 | 0.076752 | 0.765267 |
| R_PHT | 759 | -0.358675 | 0.061059 | 1.064206 |
| R_STV | 894 | -0.319639 | 0.061587 | 0.923461 |
| R_MST | 385 | -0.156272 | 0.108016 | 0.136374 |
| R_TE1p | 1157 | -0.14991 | 0.04818 | 1.301607 |
| R_TPOJ3 | 532 | -0.148761 | 0.078149 | 0.235596 |
| R_FST | 401 | -0.106974 | 0.101519 | 0.103457 |
| R_IP1 | 1019 | -0.078674 | 0.051782 | 0.443907 |
| R_TF | 1370 | -0.035853 | 0.048529 | 0.295859 |
| R_FEF | 850 | -0.029756 | 0.047562 | 0.284671 |
| R_MIP | 815 | -0.002675 | 0.047261 | 0.026925 |

## Footnotes

1 We defined human V4 as described by Winawer and Witthoft (2015).

2 We examined this because human V4 appears atypical in Figures 3A and 4A, consistent with prior literature (e.g., Goddard et al., 2011; Hansen et al., 2007; Wade et al., 2008; Winawer et al., 2010).

3 For an ROI pair to be considered significant, the binomial *p*-value associated with the mean SED across subjects had to survive FDR correction (Benjamini & Hochberg, 1995; *q* < 0.05), and the cross-species SED had to exceed chance for every individual subject. We used this conservative criterion to emphasize effects that were robust across subjects and to keep the figure sufficiently uncluttered for presentation in the manuscript. Full results are available on the website.

4 Although the contrast of words> consonant strings did not yield significant ROIs for all human subjects, a cross-species searchlight using only these human subjects/seeds identified the same macaque brain region.

5 We used the transform learned from simulated movie data using canonical HRFs (like that depicted in Figure 1D, main text, but with 2.5s TR).

